# Mitotic Regulators and the SHP2-MAPK Pathway Promote Insulin Receptor Endocytosis and Feedback Regulation of Insulin Signaling

**DOI:** 10.1101/419911

**Authors:** Eunhee Choi, Sotaro Kikuchi, Haishan Gao, Karolina Brodzik, Ibrahim Nassour, Adam Yopp, Amit G. Singal, Hao Zhu, Hongtao Yu

**Affiliations:** Howard Hughes Medical Institute, Department of Pharmacology, University of Texas Southwestern Medical Center, 6001 Forest Park Road, Dallas, TX 75390, USA; Children’s Research Institute, Departments of Pediatrics and Internal Medicine, Center for Regenerative Science and Medicine, University of Texas Southwestern Medical Center, 6000 Harry Hines Boulevard, Dallas, TX 75390, USA

## Abstract

Insulin controls glucose homeostasis and cell growth through bifurcated signaling pathways. Dysregulation of insulin signaling is linked to diabetes and cancer. The spindle checkpoint controls the fidelity of chromosome segregation during mitosis. Here, we show that insulin receptor substrate 1 and 2 (IRS1/2) cooperate with spindle checkpoint proteins to promote insulin receptor (IR) endocytosis through recruiting the clathrin adaptor complex AP2 to IR. A phosphorylation switch of IRS1/2 orchestrated by extracellularly regulated kinase 1 and 2 (ERK1/2) and Src homology phosphatase 2 (SHP2) ensures selective internalization of activated IR. SHP2 inhibition blocks this feedback regulation and growth-promoting IR signaling, prolongs insulin action on metabolism, and improves insulin sensitivity in mice. We propose that mitotic regulators and SHP2 promote feedback inhibition of IR, thereby limiting the duration of insulin signaling. Targeting this feedback inhibition can improve insulin sensitivity.

## Introduction

The pancreatic hormone insulin controls glucose homeostasis and promotes cell growth and proliferation. Dysregulation of insulin signaling is linked to human metabolic syndromes and cancer^1^. Insulin binds to the insulin receptor (IR) on the plasma membrane (PM), and triggers phosphorylation-mediated activation of crucial enzymes that regulate glucose and lipid metabolism, and cell growth and division^2^-^4^. Activated IR phosphorylates itself and the insulin receptor substrate (IRS) proteins on tyrosines. Phosphorylated IRS proteins bind to multiple downstream effectors and adaptors, and activate two major branches of insulin signaling: the phosphatidylinositol 3-kinase (PI3K)-protein kinase B (AKT) and mitogen-activated protein kinase (MAPK) pathways. The PI3K-AKT pathway mainly governs metabolic homeostasis, whereas the MAPK pathway controls cell growth and proliferation. Src homology phosphatase 2 (SHP2, also known as PTPN11) binds to the C-terminal phospho-tyrosine sites of IRS1/2 and promotes the activation of the MAPK pathway^5,6^. Mutations of IR cause severe inherited insulin resistance syndromes^7^, but the molecular mechanisms underlying insulin resistance in type 2 diabetes are complex and multifactorial^1^. One common theme is that insulin resistance in diabetic animals or patients causes ectopic accumulation of diacylglycerol and abnormal activation of novel protein kinase Cs (PKCs), which suppress insulin signaling at the level of IRS1 and possibly IR^1,8^.

The spindle checkpoint monitors kinetochore-microtubule attachment during mitosis and prevents chromosome missegregation^9-13^. In response to unattached kinetochores, the mitosis arrest deficiency 2 (MAD2) and budding uninhibited by benomyl 1 related 1 (BUBR1) proteins, as subunits of the mitotic checkpoint complex (MCC), inhibit the anaphase-promoting complex/cyclosome (APC/C) bound to its mitotic activator, the cell division cycle 20 (CDC20) protein, to delay chromosome segregation^14-19^. When all kinetochores are properly attached by microtubules, the MAD2-binding protein p31^comet^ (also called MAD2L1BP) prevents the conformational activation of MAD2^20,21^ and collaborates with the ATPase TRIP13 to disassemble MCC, thus promoting chromosome segregation^22-27^.

We have recently discovered a critical role of MAD2, BUBR1, and p31^comet^ in insulin signaling during interphase (Fig. 1a)^28^. Specifically, MAD2 and BUBR1 are required for clathrin-mediated endocytosis of IR. MAD2 directly binds to the C-terminal MAD2-interacting motif (MIM) of IR, and recruits the clathrin adaptor AP2 to IR through BUBR1. p31^comet^ prevents spontaneous IR endocytosis through blocking the interaction of BUBR1-AP2 with IR-bound MAD2. Adult liver-specific p31^comet^ knockout mice exhibit premature IR endocytosis in the liver and whole-body insulin resistance. Conversely, BUBR1 deficiency delays IR endocytosis and enhances insulin sensitivity in mice. These findings implicate dysregulation of IR endocytosis as a potential mechanism of insulin resistance.

**Fig. 1.**
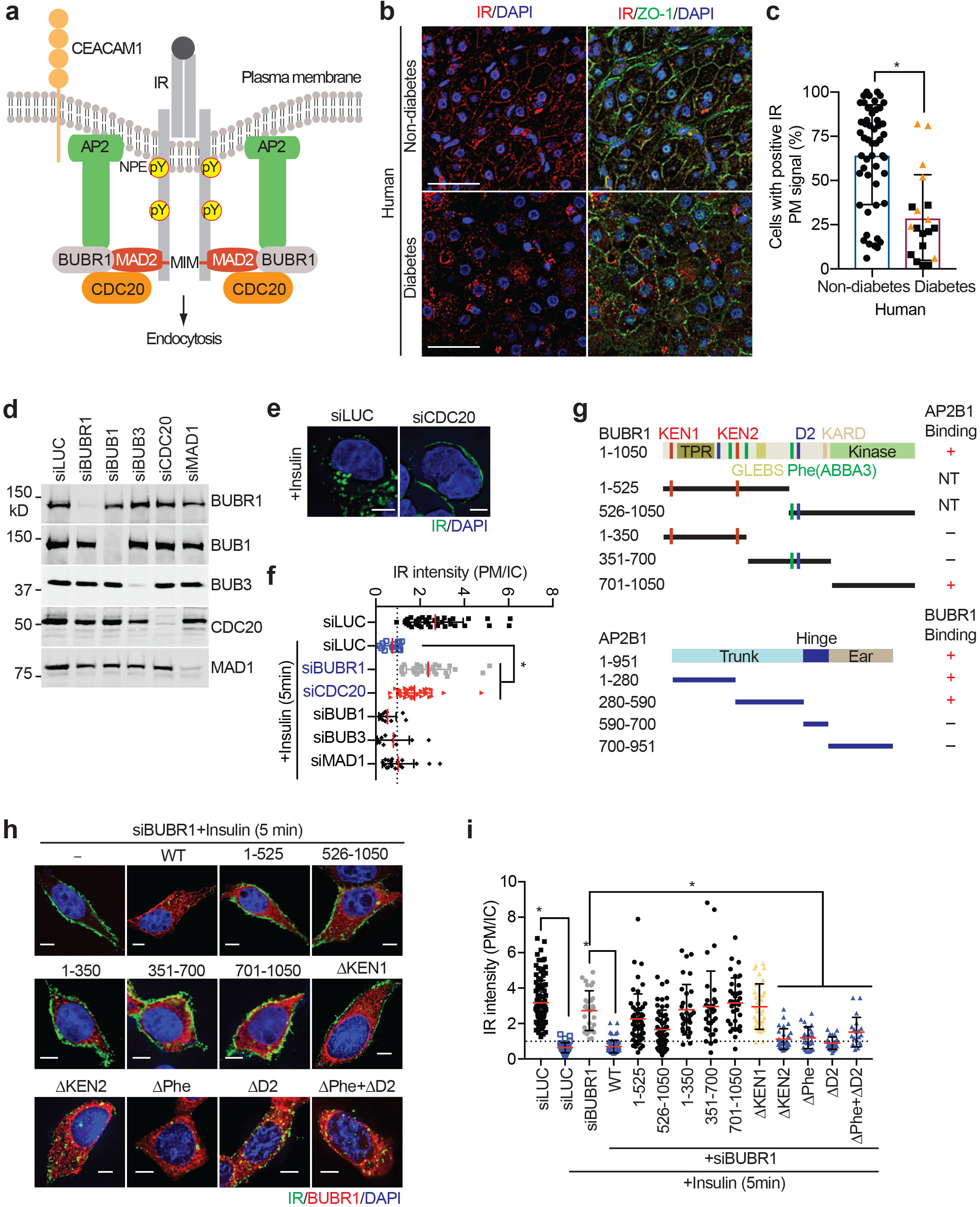
The MAD2-CDC20-BUBR1 module is required for insulin-activated IR endocytosis. **a** Model of the regulation of IR endocytosis by spindle checkpoint proteins. MIM, MAD2 interacting motif. **b** Representative images of liver specimens from human non-diabetic and diabetic patients stained with DAPI (blue) and anti-IR (red) and anti-ZO-1 (green) antibodies. Scale bars, 40 µm. **c** Quantification of the percentage of cells with positive IR PM signals in liver specimens in (**a**) (mean ± SD; *p<0.0001). The liver specimens from insulin-treated patients were presented as orange triangles. **d** Western blot analysis of lysates of HepG2 cells stably expressing IR-GFP WT that had been transfected with the indicated siRNAs. **e** Cells in (**d**) were serum starved, treated without or with 100 nM insulin for 5 min, and stained with anti-GFP antibodies. Scale bar, 10 µm. **f** Quantification of the ratios of PM and IC IR-GFP signals of cells in (**e**) (mean ± SD; *p<0.0001). **g** Domains and motifs of human BUBR1 and AP2B1. TPR, tetratricopeptide repeat domain; GLEBS, GLE2-binding sequence; KARD, kinetochore attachment regulatory domain. Fragments that can or cannot bind to AP2B1 or BUBR1 are presented as + or -, respectively. NT, not tested. The KEN1/2 boxes, destruction boxes (D1/2), and Phe boxes (also called ABBA boxes; 1/2/3) are shown as red, blue, and green bars, respectively. **h** 293FT cells stably expressing IR-GFP WT were transfected with the indicated siRNAs and Myc-BUBR1, serum starved, treated without or with 100 nM insulin for 5 min, and stained with anti-GFP (IR; green), anti-Myc (BUBR1; red), and DAPI (blue). Scale, 10 µm. **i** Quantification of the ratios of PM and IC IR-GFP signals of cells in (**h**) (mean ± SD; *p<0.0001).

In this study, we show that IRS1/2 cooperate with several spindle checkpoint proteins to promote insulin-stimulated IR endocytosis. In addition to MAD2 and BUBR1, CDC20 is also required for IR endocytosis. We further identify novel feedback regulation of IR endocytosis through a phosphorylation switch on IRS1/2 that is dependent on SHP2 and the MAPK pathway. Finally, we present evidence to suggest that targeting SHP2 might be a viable strategy to increase insulin sensitivity and treat diabetes.

## Results

### Decreased IR plasma membrane localization in human diabetics

Previous biochemical studies have shown that insulin binding to liver plasma membranes is reduced in obese and diabetic mice and humans^29,30^, suggesting that the level of insulin receptor at the plasma membrane (PM) might be reduced in diabetics. To confirm these findings, we examined the IR PM levels in liver samples from human patients using immunofluorescence staining. Because of the challenges of collecting liver sections from normal healthy individuals, we used surgical resection samples from patients with hepatocellular carcinoma that contained normal (non-malignant) and malignant tissues, and analyzed only normal hepatocytes. We performed immunofluorescence with anti-IR and anti-ZO1 (as a PM marker) antibodies on 51 non-diabetic and 19 type 2 diabetes patient samples, and analyzed IR PM levels. IR PM signals in the liver sections from type 2 diabetes patients were significantly weaker than those in non-diabetic patients (Fig. 1b,c). There was no correlation between insulin treatment and IR PM levels. These findings suggest that reduced IR PM levels might be a contributing factor to insulin resistance in human patients.

We note, however, that the reduction of IR on the cell surface in liver samples of human type 2 diabetes patients can be a consequence of insulin resistance, rather than the cause. It is also possible that the reduced IR localization at the plasma membrane is due to a decrease in overall IR expression. Despite these caveats, our immunofluorescence results are consistent with earlier findings, and suggest that mechanisms regulating IR plasma membrane levels, including endocytosis, might be defective in diabetics.

### CDC20 is required for proper IR endocytosis

We have previously shown that MAD2 and BUBR1 promote IR endocytosis through recruiting AP2, and p31^comet^ inhibits this process through antagonizing the MAD2–BUBR1 interaction (Fig. 1a)^28^. We further tested whether other well-known mitotic regulators also have a role in IR endocytosis. Depletion of CDC20, but not BUB1, BUB3, or MAD1, delayed insulin-activated IR endocytosis (Fig. 1d-f). Thus, CDC20 is required for IR endocytosis.

CDC20 binds to both MAD2 and BUBR1 and nucleates the formation of MCC that consists of BUBR1, BUB3, MAD2, and CDC20^22,31^. BUB3 is not absolutely required for the integrity and function MCC, but enhances the efficiency of MCC formation by targeting BUBR1 to kinetochores. Structures of MCC nicely explain the binding cooperativity among MAD2, CDC20, and BUBR1, and highlight the importance of the BUBR1 KEN1 motif in MCC assembly^22^. The assembly of an MCC-like complex on IR might strengthen the weak MAD2–BUBR1 interaction and help to recruit AP2 to IR.

To test this hypothesis, we first examined the region of BUBR1 responsible for IR endocytosis. The C-terminal domain of BUBR1 binds to the N-terminal trunk domain of AP2B1 (Fig. 1g and Supplementary Fig. 1a,b), consistent with a previous report^32^. Expression of wild-type (WT) BUBR1, but none of BUBR1 truncation mutants, restored IR endocytosis in cells depleted of BUBR1 (Fig. 1h,i and Supplementary Fig. 1c). BUBR1 interacts with CDC20 through multiple binding motifs^22,33-35^. We constructed BUBR1 mutants targeting KEN1 (ΔKEN1) or KEN2 (ΔKEN2) boxes in the N-terminal region, and Phe (ΔPhe, also known as ABBA3) and D boxes (ΔD2) in the middle region (Fig. 1g). The ΔKEN2, ΔPhe, ΔD2 and ΔPhe+ΔD2 restored insulin-activated IR endocytosis in cells depleted of BUBR1 (Fig. 1h,i and Supplementary Fig. 1c). However, BUBR1 ΔKEN1 could not restore IR endocytosis in BUBR1-depleted cells. Given the critical role of BUBR1 KEN1 in MCC assembly, these data suggest that IR uses an MCC-like complex to recruit AP2, except that in this context MAD2 binds to the MIM on IR, not the MIM on CDC20.

### IRS1/2 promote IR endocytosis

Insulin binding to IR promotes clathrin-mediated endocytosis of the insulin-IR complex, which regulates the intensity and duration of signaling. Two sequence motifs in IR––the NPEY^960^ and di-leucine (L^986^L) motifs––have previously been implicated in AP2 binding and IR endocytosis^36-39^ (Fig. 2a). We have shown that the MAD2-interacting motif (MIM, R^1333^ILTL) of IR binds to MAD2, which in turn recruits AP2 with the help of BUBR1 and CDC20.

**Fig. 2.**
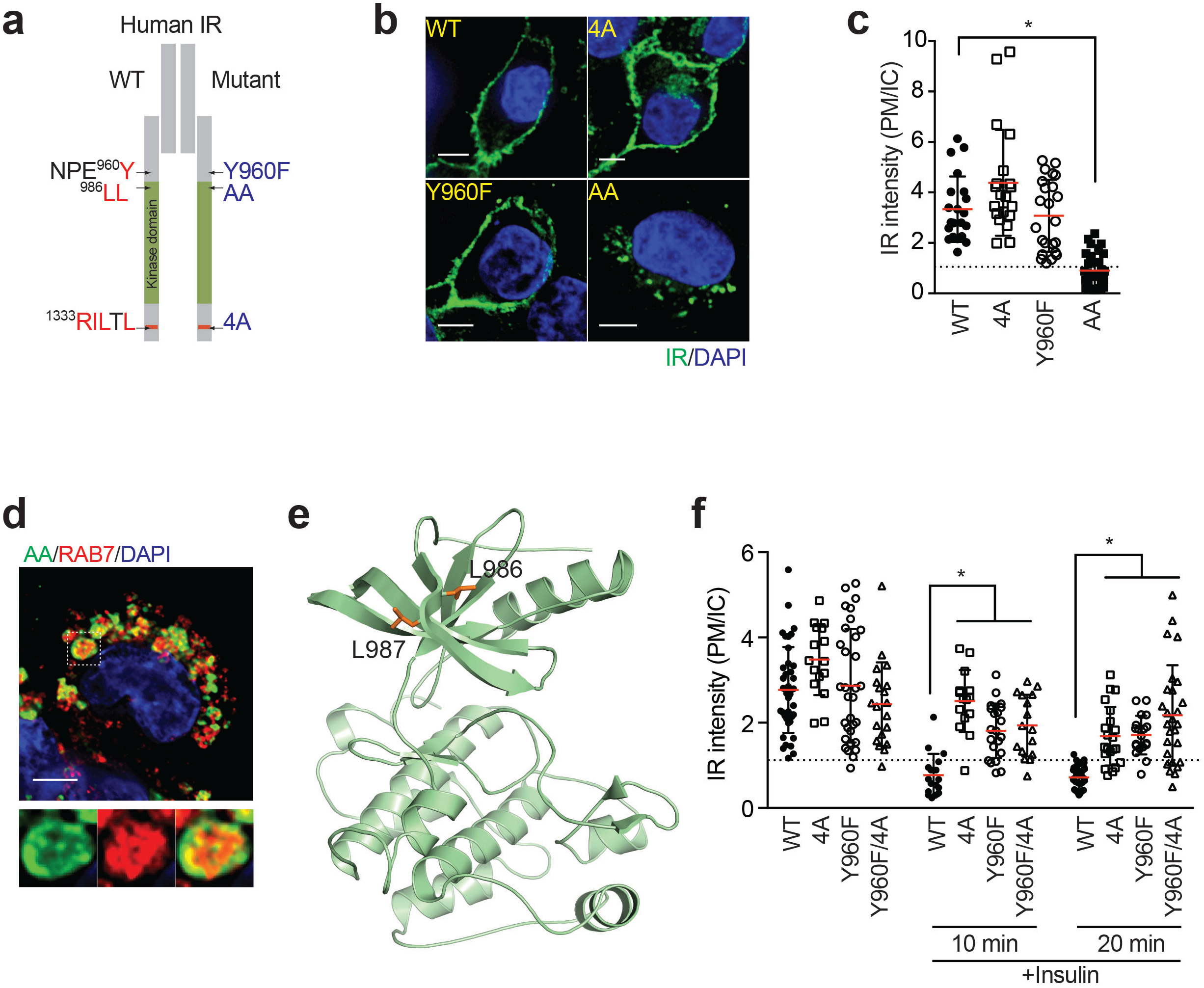
The NPEY motif and MIM of IR are required for endocytosis. **a** Schematic illustration of sequence motifs (left) and mutants (right) of IRβ. **b** HepG2 cells stably expressing IR-GFP WT, 4A, Y960F, or AA were serum starved and stained with anti-GFP antibodies. Scale bar, 10 µm. **c** Quantification of the ratios of plasma membrane (PM) and intracellular (IC) IR-GFP signals of cells in (**b**) (mean ± SD; *p<0.0001). **d** HepG2 cells stably expressing IR-GFP AA were serum starved and stained with anti-GFP and anti-RAB7 antibodies. The boxed region was magnified and shown on the right. Scale bar, 10 µm. **e** Ribbon diagram of the crystal structure of the IR kinase domain, with L986 and L987 shown as sticks. **f** HepG2 cells stably expressing IR-GFP WT, 4A, Y960F, or Y960F/4A were serum starved, treated without or with 100 nM insulin for the indicated durations, and stained with anti-GFP antibodies. Quantification of the ratios of PM and IC IR-GFP signals of cells was shown (mean ± SD; *p<0.0001).

To study the relative contributions of these IR motifs to endocytosis, we generated HepG2 cell lines stably expressing IR-GFP WT, the MIM mutant (4A), Y960F, or L986A/L987A (AA), and examined the subcellular localization of these IR-GFP proteins (Fig. 2b,c). Without insulin treatment, IR WT, 4A, and Y960F localized to the plasma membrane (PM), but IR AA was enriched in intracellular compartment (IC). A large fraction of IR AA co-localized with the late endosome marker RAB7, indicating that IR AA underwent unscheduled endocytosis and accumulated in late endosomes (Fig. 2d). Thus, the di-leucine motif is not required for IR endocytosis and actually prevents it through unknown mechanisms. The di-leucine motif is located in a β strand of the N-terminal lobe of the IR kinase domain (Fig. 2e). Mutation of this motif is expected to alter the structural integrity or activity of the IR kinase domain.

IR Y960F, 4A, and Y960F/4A mutants were less efficiently internalized after insulin stimulation (Fig. 2f). The IR Y960F/4A double mutant was not significantly more defective than the single mutants (Fig. 2f). As Y960 is phosphorylated in the activated IR^40^, defective endocytosis of IR Y960F suggests that phosphorylation of Y960 (pY960) might be required for timely IR internalization.

The phosphotyrosine-binding (PTB) domain of IRS1/2 directly binds to phosphorylated NPEY^960^ motif in activated IR^41-45^. Co-depletion of IRS1/2 blocked IR endocytosis induced by insulin, whereas depletion of either had no effect (Fig. 3a,b and Supplementary Fig. 2a). Expression of RNAi-resistant IRS1 restored IR endocytosis in cells depleted of both IRS1/2. Thus, IRS1/2 act redundantly to promote IR endocytosis likely through binding to the phospho-NPEY^960^ motif. These results can explain why the activated IR is preferentially internalized.

**Fig. 3.**
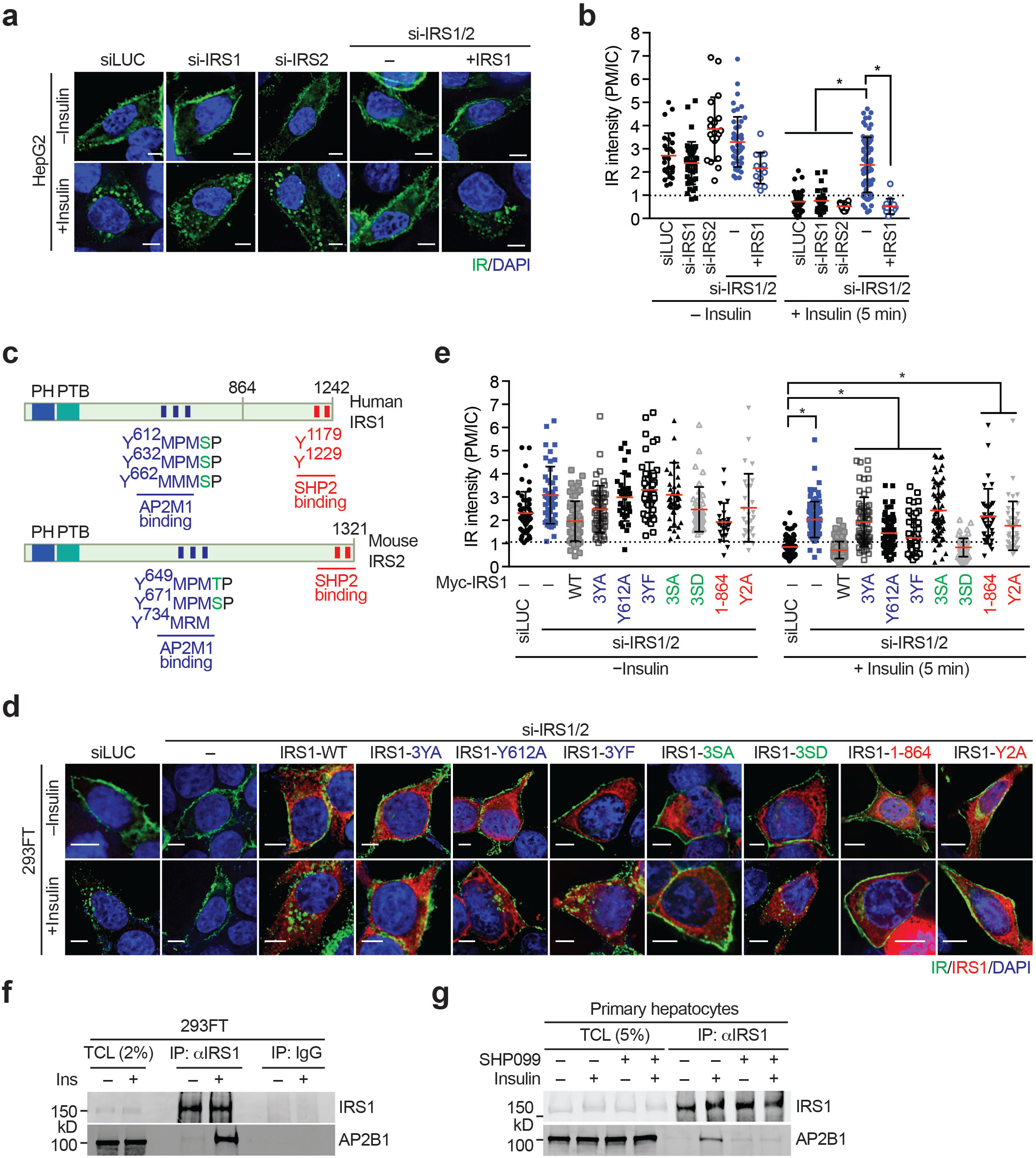
IRS1/2 are required for insulin-activated IR endocytosis. **a** HepG2 cells stably expressing IR-GFP WT were transfected with the indicated siRNAs or siRNA-resistant Myc-IRS1, serum starved, treated without or with 100 nM insulin for 5 min, and stained with anti-GFP antibodies. **b** Quantification of the ratios of PM and IC IR-GFP signals of cells in (**a**) (mean ± SD; *p<0.0001). **c** Domains and YXXΦ motifs of human IRS1 and mouse IRS2. PH, pleckstrin homology domain; PTB, phosphotyrosine-binding domain. AP2M1- and SHP2-binding regions are indicated. YXXΦ motifs and phosphotyrosine sites of IR for SHP2 binding are shown as blue and red bars, respectively. The MAPK phosphorylation sites are labeled as green letters in the sequences. **d** 293FT cells stably expressing IR-GFP WT were transfected with the indicated siRNAs or siRNA-resistant Myc-IRS1, serum starved, treated without or with 100 nM insulin for 5 min, and stained with anti-GFP (IR; green), anti-Myc (IRS1; red), and DAPI (blue). (3YA, Y612A/Y632A/Y662A; 3YF, Y612F/Y632F/Y662F; 3SA, S616A/S636A/S666A; 3SD, S616D/S636D/S666D; Y2A, Y1179A/Y1229A). **e** Quantification of the ratios of PM and IC IR-GFP signals of cells in (**d**) (mean ± SD; *p<0.0001). **f** 293FT cells were serum starved and treated without or with 100 nM insulin for 5 min. Total cell lysate (TCL), anti-IRS1 IP, and IgG IP were blotted with anti-IRS1 and anti-AP2B1 antibodies. **g** Serum-starved primary hepatocytes were treated with DMSO or 10 µM SHP099 for 2 h and treated with 100 nM insulin for 5 min. Total cell lysate (TCL), anti-IRS1 IP were blotted with anti-IRS1 and anti-AP2B1 antibodies.

IRS1 interacts with the adaptor related protein complex 1 Mu 1 subunit (AP1M1) and AP2M1 through multiple YXXΦ (X, any amino acids; Φ, bulky hydrophobic residues) motifs^46,47^. The IRS1-AP2M1 interaction negatively regulates endocytosis of insulin-like growth factor 1 receptor (IGF1R)^47^. We confirmed that in vitro translated Myc-IRS1 full-length and the YXXΦ-containing central region (residues 449-679) bound to GST-AP2M1 (Fig. 3c and Supplementary Fig. 2b-e). IRS2 is highly homologous to IRS1 and also has conserved YXXΦ motifs (Fig. 3c and Supplementary Fig. 2e)^3^. AP2M1 binds to YXXΦ motifs and promotes clathrin-mediated endocytosis^48^. Thus, IRS1/2 contribute to IR endocytosis through bridging an interaction between AP2 and activated IR.

We tested whether mutations of YXXΦ motifs in IRS1 and IRS2 disrupted AP2M1 binding. In vitro translated IRS1 (residues 449-864) bound to GST-AP2M1. Single YA mutant weakened the IRS1-AP2M1 interaction, and 3YA (Y612A/Y632A/Y662A) further reduced it (Supplementary Fig. 3a). An in vitro translated IRS2 fragment (residues 520-888) containing 7 putative YXXΦ motifs also bound to GST-AP2M1, and mutations of 4 such YXXΦ motifs (Y649A/Y671A/Y734A/Y758A) greatly reduced IRS2 binding to AP2M1 (Supplementary Fig. 3b). These results suggest that IRS1 and IRS2 directly bind to AP2M1 through multiple YXXΦ motifs.

RNAi-resistant IRS1 Y612A and 3YA mutants could not restore IR endocytosis in 293FT or HepG2 cells depleted of IRS1/2 (Fig. 3d,e and Supplementary Fig. 3d,e). Failure of these mutants to functionally complement indicates that the IRS1/2-AP2M1 interaction is required for insulin-activated IR endocytosis. We could not detect IRS1 in late endosomes during IR endocytosis, suggesting that IRS1 might dissociate from the IR complex during the uncoating of the clathrin coat and prior to the fusion of the endocytic vesicles with the endosome. Endogenous IRS1 interacted with the AP2 complex in 293FT cells and in primary mouse hepatocytes stimulated with insulin, but not in untreated cells (Fig. 3f,g). Thus, IRS1/2 bind to AP2 through canonical YXXΦ motifs in vitro and in mammalian cells, and promote insulin-activated IR endocytosis.

### The SHP2-MAPK pathway promotes IR endocytosis through IRS1/2 regulation

The tyrosine residues in YXXΦ motifs on IRS1/2 can be phosphorylated by the activated IR and be dephosphorylated by the tyrosine phosphatase SHP2^2,49^. Strikingly, the serine residues immediately following the YXXΦ motifs are well conserved (Fig. 3c and Supplementary Fig. 2e), and can be phosphorylated by ERK1/2^50-52^. ERK1/2-dependent phosphorylation has been proposed to reduce IRS1 tyrosine phosphorylation through negative feedback^50-52^, but the mechanism and function of this feedback regulation are unknown. We hypothesized that the MAPK pathway and SHP2 might regulate the IRS1/2-AP2 interaction and IR endocytosis through modulating IRS1/2 phosphorylation patterns.

To test this hypothesis, we examined the effects of inhibiting SHP2 or the MAPK pathway on insulin-activated IR endocytosis (Fig. 4a,b). The IR inhibitor (BMS536924) expectedly blocked IR endocytosis. Strikingly, the MEK inhibitors (U0126 and PD0325901) and the SHP2 inhibitor (SHP099) also inhibited IR endocytosis. By contrast, the AKT inhibitor (AKTi, VIII) did not affect IR endocytosis, indicating a specific requirement for the MAPK pathway and SHP2. Inhibitors of MPS1 (Reversine) and PLK1 (BI2546) did not appreciably inhibit IR endocytosis, ruling out the involvement of these mitotic kinases in this process.

**Fig. 4.**
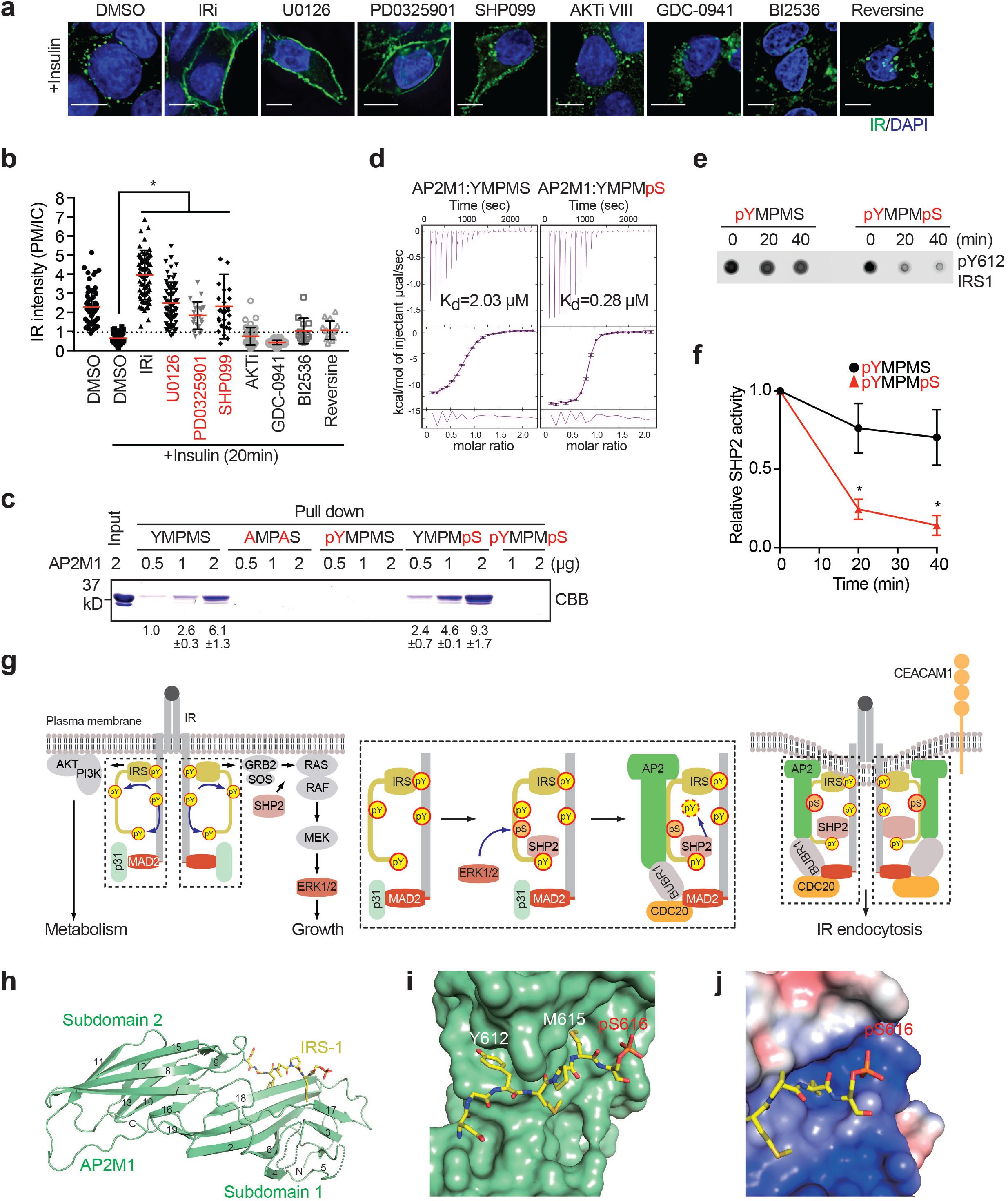
The SHP2-MAPK pathway promotes insulin-activated IR endocytosis. **a** HepG2 cells expressing IR-GFP WT were starved, treated with the indicated inhibitors for 2 h, treated without or with 100 nM insulin for 20 min, and stained with anti-GFP (IR; green) and DAPI (blue). **b** Quantification of the ratios of PM and IC IR-GFP signals of cells in (A) (mean ± SD; *p<0.0001). **c** Binding of IRS1 peptides to AP2M1 (residues 160-435). Input and proteins bound to IRS1-peptide beads were analyzed by SDS-PAGE and stained with Coomassie (CBB). The relative band intensities are shown below (mean ± SD; n=4 independent experiments). **d** Isothermal titration calorimetry (ITC) analysis of binding between IRS1 peptides and AP2M1 (residue 160-435), with K_d_ indicated. **e** The IRS1 peptides were incubated with active SHP2 for the indicated durations, spotted onto membranes, and detected with the anti-pY612-IRS1 antibody. **f** Quantification of the relative SHP2 activity in (**e**) (mean ± SD; n=4 independent experiments; *p<0.0001). **g** Model of the regulation of insulin-activated IR endocytosis by a phosphorylation switch on IRS1/2. Insulin-bound IR phosphorylates itself and IRS1/2, and activates the PI3K-AKT and MAPK pathways. SHP2 acts upstream of RAS-RAF and promotes the activation of MAPK pathway. p31^comet^ binds to the IR-bound MAD2 and blocks IR-AP2 association to prevent premature IR endocytosis. In feedback regulation, activated ERK1/2 phosphorylate S616 and other sites on IRS1. SHP2 binds to the C-terminal phospho-tyrosine site on IRS1 and dephosphorylates pY612 of the doubly phosphorylated IRS1 (pY612/pS616), thus promoting IRS1-AP2M1 association. p31^comet^ is released from MAD2 by an unknown mechanism, allowing the assembly of an MCC-like complex on IR. MAD2- and IRS1/2-dependent AP2 recruitment and clustering trigger clathrin-mediated IR endocytosis. **h** Ribbon diagram of the crystal structure of AP2M1 (residues 160-435) bound to pS-IRS1. pS-IRS1 is shown as sticks. **i** Surface drawing of AP2M1, with pS-IRS1 shown as sticks. **j** A close-up view of the surface drawing of AP2M1 colored by its electrostatic potential (blue, positive; red, negative; white, neutral). pS-IRS1 is shown as sticks.

The tyrosine residues in YXXΦ motifs on IRS1/2 also bind PI3K^53,54^. It is formally possible that the failure of IRS1 3YA mutant to restore IR endocytosis in cells depleted of IRS1/2 was an indirect consequence of reduced PI3K activity. To test this possibility, we checked the effect of IRS1 3YA on insulin signaling (Supplementary Fig. 3e). IRS1/2 depletion was sufficient to inhibit IR endocytosis, but was insufficient to inhibit insulin signaling at insulin concentrations used in our IR endocytosis assays. Expression of IRS1 WT or the 3YA mutant did not appreciably alter insulin signaling. Furthermore, the PI3K inhibitor (GDC-0941) did not affect insulin-activated IR endocytosis (Fig. 4a,b). This result argues against reduced PI3K activity as the underlying reason for the observed defects of insulin-activated IR endocytosis in IRS1/2-depleted cells, and further confirms that the PI3K-AKT pathway is not involved in this process.

We next chemically synthesized the unphosphorylated and phosphorylated IRS1 peptides containing ^612^YMPMS and examined their binding to AP2M1 (Fig. 4c). The unphosphorylated IRS1 peptide (YMPMS) bound to AP2M1, but the mutant peptide with Y612 and M615 replaced by alanine (AMPAS) did not. Phosphorylation of the serine in the YMPMS motif (YMPMpS) enhanced AP2M1 binding. Isothermal titration calorimetry (ITC) measurements confirmed that the phospho-serine IRS1 peptide (pS-IRS1) indeed bound to AP2M1 with higher affinity (K_d_ = 280 nM), as compared to the unphosphorylated peptide (K_d_ = 2.03 µM) (Fig. 4d). Thus, phosphorylation of IRS1 at S616, a known ERK1/2 site, enhances IRS1 binding to AP2M1 in vitro. Phosphorylation of the tyrosine in the YMPMS motif (pYMPMS and pYMPMpS) abolished AP2M1 binding (Fig. 4c), suggesting that tyrosine dephosphorylation of IRS1 is required for AP2M1 binding. Consistent with these in vitro findings, SHP2-binding-deficient IRS1 mutants (residues 1-864 or Y2A, Y1179A/Y1229A) could not restore insulin-activated IR endocytosis in 293FT or HepG2 cells depleted of IRS1/2 (Fig. 3d,e and Supplementary Fig. 3c,d).

Comparison of the dephosphorylation kinetics of singly (pYMPMS) or doubly (pYMPMpS) phosphorylated IRS1 peptides revealed that pS616 on IRS1 promoted pY612 dephosphorylation by SHP2 (Fig. 4e,f). Therefore, aside from directly augmenting the IRS1-AP2M1 interaction, serine phosphorylation of IRS1 indirectly promotes AP2M1 binding through enhancing the tyrosine dephosphorylation of IRS1 by SHP2 in vitro. Consistent with a role of ERK1/2-dependent phosphorylation of IRS1 in IR endocytosis, expression of the RNAi-resistant IRS1 phospho-mimicking mutant (3SD), but not the phospho-deficient mutant (3SA), restored IR endocytosis in 293FT cells depleted of IRS1/2 (Fig. 3d,e).

Taken together, our results support the following mechanism for insulin-activated IR endocytosis (Fig. 4g). The activated IR phosphorylates the tyrosine residues in YXXΦS motifs and the C-terminal SHP2-docking sites of IRS1/2, and stimulates the PI3K-AKT and MAPK pathways. In a negative feedback mechanism, activated ERK1/2 phosphorylate the serines in YXXΦS motifs on IRS1/2 and assist SHP2 to dephosphorylate IRS1/2. The IRS1/2 YXXΦS motifs with the serine phosphorylated and tyrosine dephosphorylated bind to AP2 with optimal affinities, promoting clathrin-mediated endocytosis of IR.

### Structural basis of the phospho-regulation of IR endocytosis

We next determined the crystal structure of AP2M1 (residues 160-435) bound to the serine-phosphorylated YXXΦS motif from IRS1 (pS-IRS1) (Supplementary Table 1). The overall structure of the AP2M1–pS-IRS1 complex was similar to those of previously determined structures of AP2M1 bound to YXXΦ motifs. AP2M1 contained two interlinked β-sandwich subdomains: subdomain 1 (β1-6 and 17-19) and subdomain 2 (β7-16) (Fig. 4h). The pS-IRS1 peptide binds at the edges of strands β18 and β17 in subdomain 1, and interacts with residues from strands β1, β17, and β18 (Fig. 4h,i). In particular, Y612 and M615 make extensive hydrophobic interactions with AP2M1. The RNAi-resistant IRS1 3YF mutant with tyrosines in the YXXΦS motifs replaced by phenylalanines could not fully restore IR endocytosis in 293FT cells depleted of IRS1/2 (Fig. 3d,e). The hydroxyl group of Y612 forms a hydrogen bond with D176 in β1, providing an explanation for why phenylalanines cannot functionally substitute for tyrosines. Phosphorylation of Y612 is expected to introduce both static hindrance and unfavorable electrostatic interactions with D176, explaining why tyrosine phosphorylation of YXXΦS motifs disrupts the IRS1–AP2 interaction.

We did not observe well-defined electron density for pS616 in IRS1, despite its ability to enhance the IRS1–AP2 interaction. pS616 is located in the vicinity of a positively charged patch on AP2M1 formed by residues K405, H416, and K420 (Fig. 4j), suggesting that the phosphoserine might engage in favorable electrostatic interactions with this basic patch. Mutations of H416 and K420 did not, however, reduce IRS1 binding (Supplementary Fig. 3f,g). Mutation of K405 destabilized the AP2M1 protein and reduced its binding to both the phosphorylated (pS616) and unphosphorylated IRS1 peptides. Thus, consistent with the lack of electron density, pS616 does not make defined electrostatic interactions with specific acceptor residues, and interacts with the positively charged patch as one structural entity.

### SHP2 promotes IR endocytosis in mice

Liver-specific SHP2 knockout (KO) mice show increased insulin sensitivity^55,56^, suggesting that SHP2 attenuates certain aspects of insulin signaling in the liver. We examined whether SHP2 inhibition could improve insulin sensitivity. The allosteric SHP2 inhibitor, SHP099, stabilizes the inactive conformation of SHP2, thus inhibiting its phosphatase activity^5^. Wild type mice maintained on high-fat diet (HFD) for 5 weeks were treated with SHP099 (60 mg/kg body weight) by daily oral gavage for 6 days, and then tested for glucose and insulin tolerance. Strikingly, SHP099 administration markedly increased glucose tolerance and insulin sensitivity in HFD-fed mice (Fig. 5a,b). SHP099 did not change the body weight of mice fed HFD (Fig. 5c). Insulin stimulation caused IR endocytosis and reduced the IR staining at the PM in mouse liver sections (Fig. 5d,e). SHP099 delayed insulin-activated IR endocytosis. Finally, SHP099 inhibited the insulin-stimulated IRS1–AP2 interaction in primary hepatocytes (Fig. 3g).

**Fig. 5.**
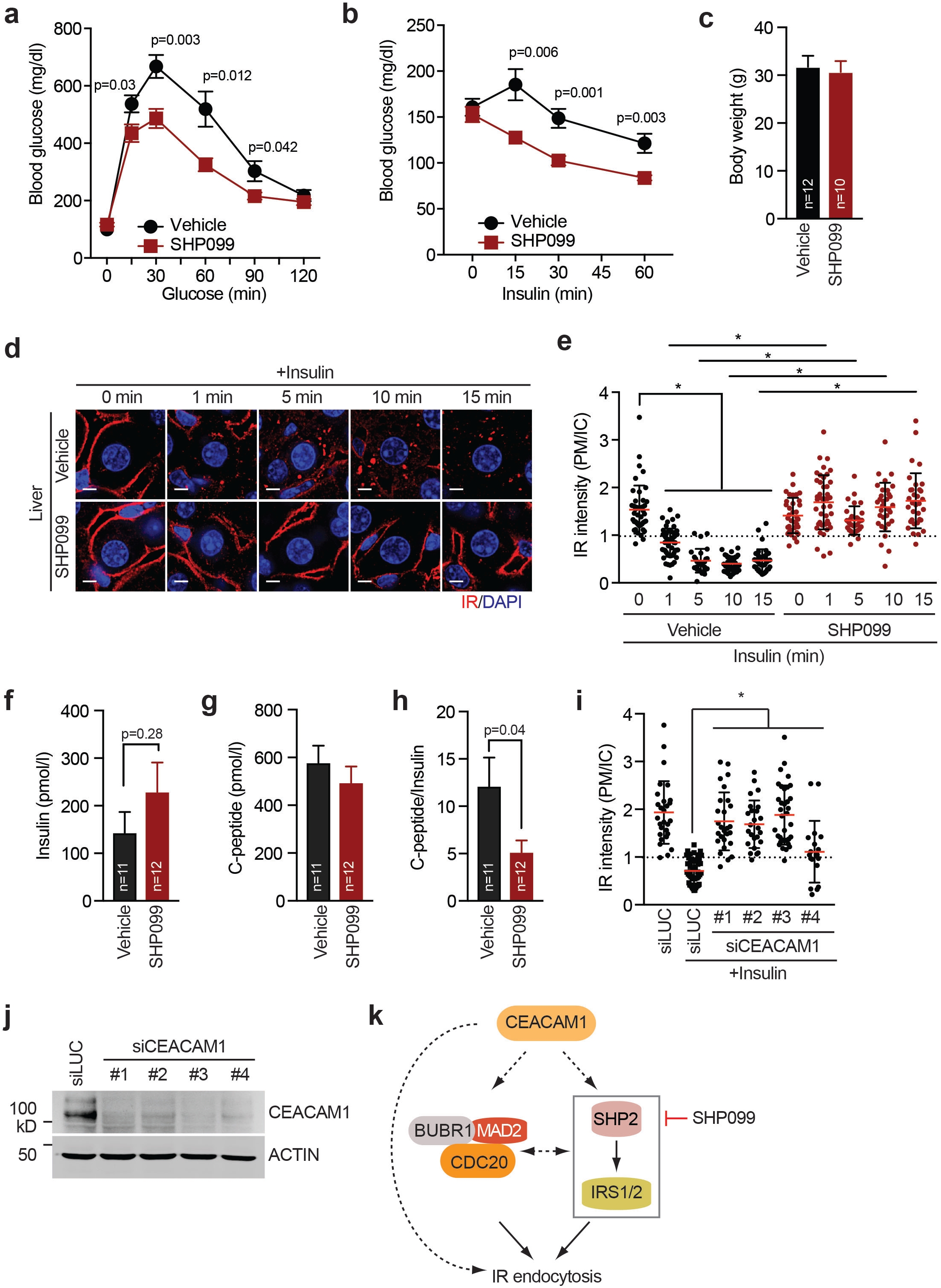
SHP2 inhibition delays IR endocytosis and improves insulin sensitivity in mice. **a**,**b** Glucose tolerance test (**a**) and insulin tolerance test (**b**) of male mice fed HFD for 5 weeks. The mice were administered vehicle or SHP099 for 6 days. At 1 day after the last drug administration, experiments were performed. Vehicle, n=12; SHP099, n=10; mean ± SEM. **c** Body weight of mice administered vehicle or SHP099 at 7 days post administration. Mean ± SD. **d** HFD-fed mice were administered vehicle or SHP099 for 5 days. The mice were fasted overnight and administered vehicle or SHP099 once more. At 2 h after the last administration, the mice were injected with or without 1 U insulin via inferior vena cava. The livers were collected at the indicated time points and the sections were stained with anti-IR (red) and DAPI (blue). Scale bars, 5 µm. **e** Quantification of the ratios of plasma membrane (PM) and intracellular compartments (IC) IR signals of the livers in (**d**) (mean ± SD; *p<0.0001). **f**-**h** The levels of fasting serum insulin (**f**) and C-peptide (**g**), and the ratio of C-peptide:insulin (**h**) in mice fed normal chow or HFD for 5 weeks. The mice were administered vehicle or SHP099 for 6 days. **i** HepG2 cells stably expressing IR-GFP were transfected with CEACAM1 siRNAs, serum starved, treated without or with 100 nM insulin form 5 min, and stained with anti-GFP and DAPI. Quantification of the ratios of PM and IC IR-GFP signals of cells was shown (mean ± SD). **j** Western blot analysis of cell lysates in (**i**). **k** Model of the regulation of insulin-activated IR endocytosis by CEACAM1, the MAD2–CDC20– BUBR1 module, and the SHP2-IRS1/2 module.

To further confirm the requirement of SHP2 in promoting IR endocytosis in vivo, we introduced adeno-associated viruses 8 (AAV) encoding control (Ctrl) or SHP2 short-hairpin RNAs (shRNA) into mice fed HFD via tail-vein injection. The SHP2 protein level in the liver from mice treated with AAV-SHP2 shRNA was reduced by ∼70% as compared to that in mice treated with Ctrl shRNA (Supplementary Fig. 4a). By contrast, the SHP2 protein level in WAT and skeletal muscle was not effectively depleted by SHP2 shRNA. Thus, as expected, SHP2 shRNA delivered by AAV was most effective in the liver. Consistent with the phenotypes of SHP099 administration, AAV-SHP2 shRNA treatment markedly increased glucose tolerance and insulin sensitivity in mice fed with HFD (Fig. 5a,b). AAV-SHP2 shRNA treatment did not change body weight (Fig. 5c). Importantly, AAV-mediated SHP2 silencing inhibited insulin-activated IR endocytosis in the liver (Fig. 5d,e). These results confirm a role of SHP2 in promoting IR endocytosis and in metabolic homeostasis in vivo.

Liver is a major site for insulin clearance and defects in this process can cause hyperinsulinemia^57-59^. Hepatic insulin clearance is mainly mediated by IR endocytosis, as liver-specific IR KO mice and mice deleted of the carcinoembryonic antigen-related cell adhesion molecule 1 (CEACAM1), a key regulator of IR endocytosis, develop severe hyperinsulinemia^57,59^. We thus examined the effects of SHP099 on insulin clearance. To estimate insulin clearance, we examined the levels of insulin and C-peptide, a cleavage product of proinsulin, and determined the ratio of the serum levels of C-peptide and insulin. The fasting insulin level in mice fed normal chow was not altered by SHP099 administration, despite the inhibition of IR endocytosis in the liver (Fig. 5f). In HFD-fed mice, SHP099 slightly increased the fasting insulin level. The serum levels of C-peptide were similar to those in vehicle-treated groups in both conditions (Fig. 5g), suggesting that SHP099 did not alter insulin secretion. As a result, the C-peptide:insulin ratio in mice fed HFD after SHP099 administration showed a mild reduction as compared to the control group (Fig. 5h). Therefore, despite causing defective IR endocytosis in the liver, SHP2 inhibition led to a mild defect in insulin clearance, but not severe hyperinsulinemia as observed in the CEACAM1 KO mice.

As shown in Fig. 4b, the inhibitory effect of SHP099 on IR endocytosis was not as complete as that of the IR inhibitor at 20 min after insulin stimulation, suggesting that there might be SHP2-independent IR endocytosis mechanisms. We have confirmed a requirement for CEACAM1 in IR endocytosis in our cellular assays (Fig. 5i,j). It is possible that CEACAM1 is required for all mechanisms of IR endocytosis whereas SHP2 only regulates the IRS1/2 branch (Fig. 5k). Future studies are needed to define the relationships and relative contributions of the various regulators of IR endocytosis.

### SHP2 inhibition enhances insulin-activated AKT pathway in mice

We next examined the effect of SHP2 inhibition on insulin signaling in the liver, epididymal white adipose tissue (WAT), and skeletal muscle from mice fed with HFD. We monitored the insulin-induced phosphorylation of IRS1 (pY608 and pS612; equivalent to pY612 and pS616 of human IRS1, respectively), IR (pY1152/1153 and pY962; equivalent to pY1150/1151 and pY960 of human IR, respectively), AKT (pT308), and ERK1/2 (pT202/Y204, pERK1/2). Indeed, SHP2 inhibition enhanced and prolonged the IRS1 pY608 signal in the liver, indicating that SHP2 is a relevant phosphatase for this phosphorylation in vivo (Fig. 6a,b). Consistent with a previous report ^5^, SHP099 inhibited the activation of the MAPK pathway by insulin in the liver. SHP2 inhibition attenuated insulin-induced IRS1 pS612 signal, consistent with the fact that ERK1/2 mediate this phosphorylation in vivo. By contrast, insulin-triggered activating phosphorylation of AKT was significantly increased and prolonged in the liver from SHP099-treated mice. Consistent with an inhibition of IR endocytosis, the total IR levels in both groups of mice were increased by SHP099 treatment whereas the ratios of phospho-IR to total IR were not altered. These results suggest that targeting SHP2 can block the feedback regulation of IR endocytosis by selectively inhibiting the MAPK pathway. Suppressed IR endocytosis prolongs signaling through the PI3K-AKT pathway, which regulates metabolism and does not depend on SHP2.

**Fig. 6.**
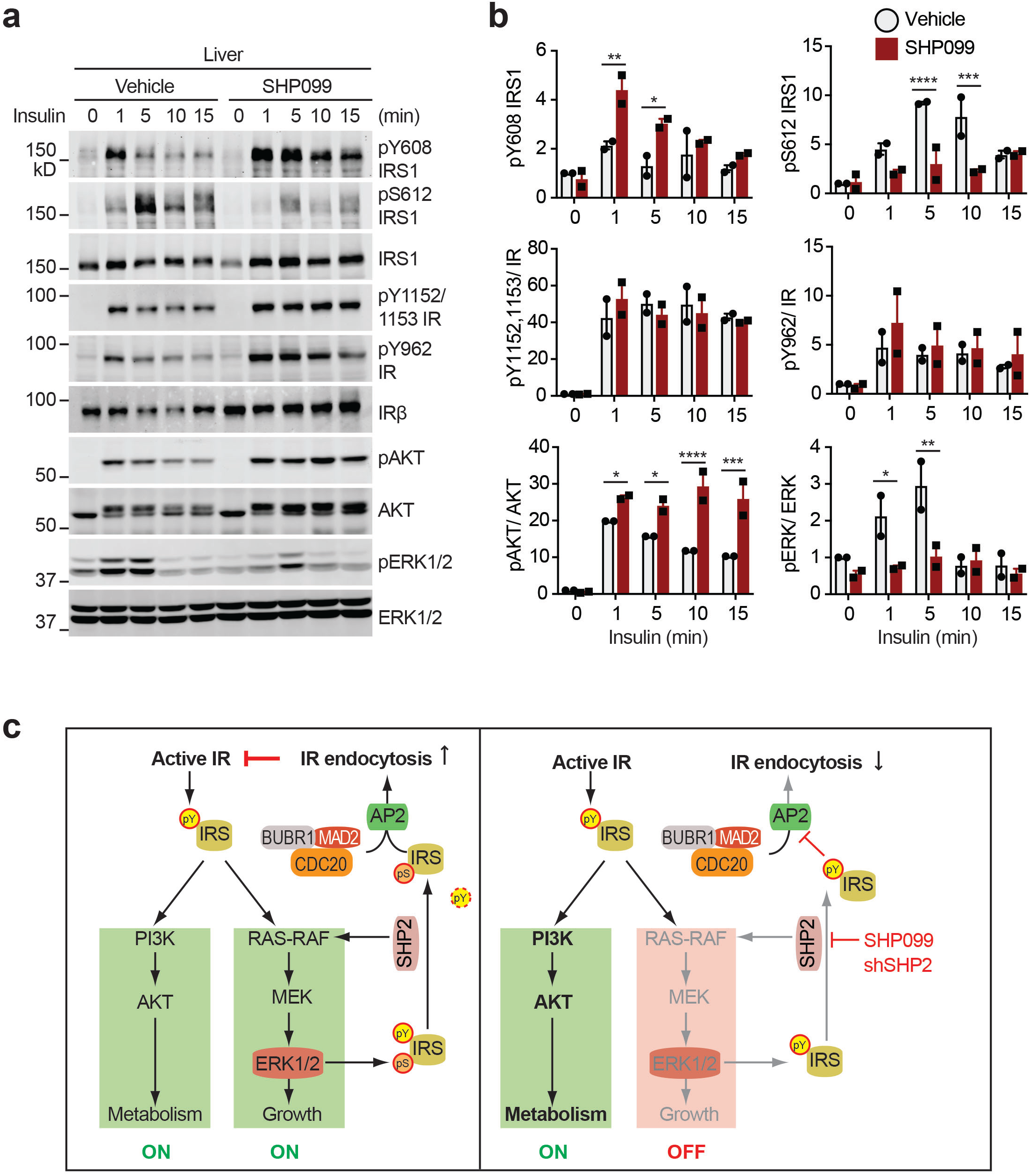
Mitotic regulators and SHP2 promote feedback inhibition of IR. **a** Insulin signaling in the liver from mice fed HFD for 5 weeks. The mice were administered vehicle or SHP099 for 5 days, fasted overnight, and administered vehicle or SHP099 once more. At 2 h after the last administration, the mice were injected with or without 1 U insulin via inferior vena cava. The livers were collected at the indicated time points. Lysates were prepared from these tissues and subjected to quantitative immunoblotting with the indicated antibodies. **b** Quantification of the blots in (**a**). Mean ± SD; *p<0.05, **p<0.01, ***p<0.001, and ****p<0.0001. **c** Targeting feedback regulation of IR endocytosis for diabetes treatment. Left panel depicts the feedback regulation of IR endocytosis by ERK1/2 and SHP2 during unperturbed insulin signaling. Right panel illustrates the mechanism by which SHP2 inhibitor (SHP099) or shRNA blocks growth-promoting IR signaling and IR endocytosis, and prolongs insulin signaling through the PI3K-AKT pathway, which controls metabolism.

SHP099 greatly inhibited the activation of the MAPK pathway and the serine phosphorylation on IRS1 (pS612) upon insulin stimulation in skeletal muscle (Supplementary Fig. 5a,b). However, the IRS1 pY608 level in skeletal muscle from SHP099-treaed mice exhibited only a slight increase, and the levels of activating phosphorylation of AKT were similar to those in vehicle-treated mice. SHP099 did not appreciably alter the insulin-stimulated activation of the PI3K-AKT or MAPK pathways in WAT (Supplementary Fig. 5c,d). These data suggest that SHP2 inhibition has distinct effects on insulin signaling and possibly IR endocytosis in different tissues. The underlying reasons for these variations among tissues are unknown at present. Because SHP099 only increases AKT signaling in the liver, but not in skeletal muscle or WAT, our findings suggest that the liver is a major site of action for the insulin-sensitizing effect of the SHP2 inhibitor in vivo. We cannot, however, rule out the contributions of other tissues to the observed phenotypes.

## Discussion

We have elucidated a feedback regulatory mechanism of IR endocytosis and identified SHP2 as a molecular target whose inhibition delays insulin-activated IR endocytosis (Fig. 6c). Targeting this feedback regulation of IR endocytosis prolongs the metabolic branch of insulin signaling and improves insulin sensitivity in mice.

The mechanism of IR endocytosis was extensively studied for decades. Our recent discovery that mitotic checkpoint regulators, including MAD2 and BUBR1, are required for IR endocytosis promoted us to re-examine the mechanism of IR endocytosis. In addition to the recently discovered MAD2-interacting motif (MIM), two other sequence motifs in IR, the NPXY^960^ motif and the di-leucine motif (L^986^/L^987^), had previously been implicated in AP2 binding and IR endocytosis^36-39^. In this study, we have shown that the dileucine motif is not required for IR endocytosis. The NPXY motif is indeed required for IR endocytosis. Instead of directly binding to AP2, the phosphorylated NPXY motif binds to IRS1/2, which in turn recruits AP2 through multiple YXXΦ motifs. We propose that IRS1/2 bound to pY999 of IR provide one binding site for AP2, and the MAD2–CDC20–BUBR1 complex bound to the MIM of IR provides another AP2-binding site. These two modules collaborate to recruit AP2 to IR, triggering IR endocytosis. Alternatively, as IRS1/2 contain multiple functional YXXΦ motifs, each of these motifs and MAD2–CDC20–BUBR1 may mediate the recruitment of one AP2 complex, leading to the clustering of multiple AP2 molecules on IR and efficient assembly of the clathrin coat.

Our findings presented herein implicate that an MCC-like mitotic regulator module is assembled onto IR to control its endocytosis in interphase. These results raise the intriguing possibility that IR might reciprocally regulate MCC assembly and spindle checkpoint signaling during mitosis. It will be interesting to examine the effect of the connection between IR and mitotic checkpoint regulators on aneuploidy and spindle checkpoint activities.

Endocytosis of a cell surface receptor normally occurs after the receptor has been activated and has transduced its signals downstream. The mechanism ensuring that IR endocytosis only occurs after downstream signaling is unknown. In this study, we have shown that the MAPK pathway is required for IR endocytosis. Activated ERK1/2 phosphorylates the YXXΦS motifs in IRS1/2 and promotes dephosphorylation of the tyrosine in these motifs by SHP2 in vitro and in vivo. The YXXΦS motifs containing phospho-serine and unphosphorylated tyrosine are optimal ligands for AP2. Because ERK1/2 activation is a critical downstream event of insulin signaling, the requirement for an active MAPK pathway in IR endocytosis constitutes important feedback regulation that ensures proper insulin signaling. Chemical inhibition of SHP2 is an effective way to disrupt this feedback regulation, as it not only directly prevents the removal of the negative tyrosine phosphorylation, but also inhibits the MAPK pathway and blocks the installation of the positive serine phosphorylation. Whether the MAD2-CDC20-BUBR1 module is under similar feedback regulation by the MAPK pathway is an interesting question for future studies.

Our study suggests that inhibition of SHP2 delays IR endocytosis, prolongs IR signaling through the PI3K-AKT pathway, and improves insulin sensitivity in the mouse. SHP2 inhibition prolongs phosphorylation of YXXΦ motifs on IRS1/2. Because these phospho-tyrosine motifs also interact with PI3K, the enhanced tyrosine phosphorylation of these motifs as a result of SHP2 inhibition may directly promote PI3K activation. This direct effect may contribute to the effectiveness of SHP2 inhibitors in counteracting insulin resistance. Genetic suppression of IR endocytosis in other ways, without the inhibition of SHP2, will better define the role of IR endocytosis during insulin signaling.

Future studies are required to clarify the potential role of premature IR endocytosis in the pathogenesis of human insulin resistance syndromes. Mutations of IR are known to cause inherited severe insulin resistance syndromes^7^, but the mechanisms by which these mutations affect IR function have not been systematically explored. It will be interesting to test whether these mutations cause premature IR endocytosis and whether SHP2 inhibitors can recover the IR PM levels and insulin sensitivity.

Hyperactivation of signaling through receptor tyrosine kinases (RTKs) drives cancer progression. Chemical inhibitors of RTKs are widely used in the clinic to treat human cancers. A common signaling pathway downstream of RTKs is the RAS-RAF-MAPK pathway, which promotes cell proliferation and survival. SHP2 acts upstream of RAS-RAF to activate the MAPK pathway^60,61^. Thus, intense efforts have been devoted to develop chemical inhibitors of SHP2, which are expected to have therapeutic potential in cancer chemotherapy. SHP099, a specific allosteric inhibitor of SHP2, has been shown to have efficacy in targeting RTK-driven cancers in animal models^5^. Although phenotypes of SHP2 KO mice suggest potential adverse effects of SHP2 inhibition^62-66^, SHP099 administration into mice did not show significant toxic effects in that previous study^5^ and in the present study. Our study shows that SHP2 inhibition improves systemic insulin sensitivity in mice. Thus, SHP2 inhibitors, such as SHP099, can be potentially repurposed to treat type 2 diabetes. Obesity increases the risks of both diabetes and certain types of cancers in humans^67^. The prevalence of all three conditions (obesity, diabetes, and cancer) has increased in recent years. SHP2 inhibitors may be particularly beneficial to patients who suffer from both diabetes and cancer.

## Methods

### Mice

Animal work described in this manuscript has been approved and conducted under the oversight of the University of Texas Southwestern Institutional Animal Care and Use Committee. All animals were maintained in a specific antigen-free barrier facility with 12 h light/dark cycles (6AM on and 6 PM off). Mice were fed a standard rodent chow (2016 Teklad Global 16% protein rodent diet, Harlan Laboratories). For inducing insulin resistance, C57BL/6J (Stock No. 000664, Jackson laboratory) were fed a high-fat (60%) diet (OpenSource Diets, Cat. No. D12492).

Glucose and insulin tolerance tests, and metabolic analysis were performed as described previously^28^. For in vivo pharmacological assays, 6-8-week-old male mice were fed high-fat diet (HFD) for 5 weeks. Two days before drug administration, mice were switched to normal chow. SHP099 (MedChem Express) was dissolved in DMSO and diluted into a 0.5% hypromellose and 0.1% Tween-80 solution. 60 mg/kg of SHP099 was administered by daily oral gavage for 6 days. For glucose tolerance test, mice were fasted for 14 h, and their blood glucose levels (T=0) were measured with tail bleeding using a glucometer (AlphaTRAK). Then, 2 g of glucose/kg of body weight was injected intraperitoneally. Blood glucose levels were measured at the indicated time points after glucose injection. For insulin tolerance test, mice fasted for 4 h were injected intraperitoneally with recombinant human insulin (Eli Lilly) at 1 U/kg body weight, and their blood glucose levels were measured at the indicated time points after injection.

### Reagents

Generation of rabbit polyclonal antibodies against GST, MAD1, CDC20 and BUB1 was described previously^28,68-70^. The following antibodies were purchased from commercial sources: anti-ZO-1/TJP1 and anti-ACTIN (MA137018; Thermo Scientific); anti-IR-pY1150/1151 (19H7; labeled as pY1152/1153 IR for mouse IR in this study), anti-IRS1-pS616 (C15H5; labeled as pS612 IRS1 for mouse IRS1 in this study), anti-AKT (40D4), anti-pT308 AKT (D25E6), anti-ERK1/2 (L34F12), anti-pERK1/2 (197G2), anti-SHP2 (D50F2) and anti-RAB7 (D95F2, Cell Signaling); anti-IRS1-pY612 (labeled as pY608 IRS1 for mouse IRS1 in this study), anti-IR-pY972 (labeled as pY962 IR for mouse IR in this study) and anti-IR (CT-3, Millipore); anti-AP2B1 (BD Biosciences); anti-IRS2 (EPR904) and anti-AP2M1 (EP2695Y, Abcam); anti-GFP and anti-MYC (9E10; Roche); anti-IRS1 (A301-158A, Bethyl laboratory); anti-CEACAM1 (283340, R&D Systems); anti-ACTIN (C-4), anti-IR (CT-3); and anti-mCherry (1C51, Novus).

The small interfering RNAs (siRNAs) were synthesized by Dharmacon (Lafayette, CO) and had the following sequences: human BUBR1 (GGA CAC AUU UAG AUG CAC Utt) ^28^; human CDC20 (AGA ACA GAC UGA AAG UAC UUU); human MAD1 (GAG CAG AUC CGU UCG AAG UUU)^68^; human BUB3 (GAG UGG CAG UUG AGU AUU U); human BUB1 (GAG UGA UCA CGA UUU CUA A)^71^; human IRS1 (GAA CCU GAU UGG UAU CUA C dTdT); human IRS2 (On-TARGETplus human IRS2 (8660) siRNA-SMARTpool); human CEACAM1 #1 (CCA UCA UGC UGA ACG UAA A); human CEACAM1 #2 (GAU CAU AGU CAC UGA GCU A); human CEACAM1 #3 (CGU AUU GGU GUG AGG UCU U); human CEACAM1 #4 (CCA UUA AGU ACA UGU GCC A); siLUC (UCA UUC CGG AUA CUG CGA U). The cDNAs encoding human IRS1 and human AP2M1 were purchased from Thermo Scientific. pBabe-puro mouse IRS2 was a gift from Dr. Ronald Kahn (Addgene plasmid #11371). The siRNA-resistant and YXXΦ motif mutants of IRS1 were generated by site-directed mutagenesis (Agilent Technology). IRS1 peptides (YMPMS, CHTDDGYMPMSPGVA; AMPAS, CHTDDGAMPASPGVA; pYMPMS, CHTDDGpYMPMSPGVA; YMPMpS, CHTDDGYMPMpSPGVA; pYMPMpS, CHTDDGpYMPMpSPGVA) were chemically synthesized at KareBay Biochem, Inc.

For testing the effects of kinase inhibitors on IR endocytosis, the cells were serum starved for 14 h and inhibitors were added at 2 h before insulin treatment. Inhibitors used in this study were as follows: the IR kinase inhibitor BMS536924 (2 µM; Tocris), the MEK inhibitors U0126 (40 µM; Cell signaling) and PD0325901 (10 µM; Selleck Chemicals), the SHP2 inhibitor SHP099 (10 µM; Medchem express), the AKT inhibitor VIII (5 µM, Calbiochem), the PI3K inhibitor GDC-0941 (10 µM; Selleck Chemicals), the PLK1 inhibitor BI2536 (200 nM, Selleck Chemicals), and the MPS1 inhibitor Reversine (1 µM, Sigma).

### Cell culture, transfection, and viral infection

293FT and HepG2 cells were cultured in high-glucose DMEM supplemented with 10% (v/v) FBS, 2 mM L-glutamine, and 1% penicillin/streptomycin. Plasmid transfections into 293FT and HepG2 cells were performed with Lipofectamine^™^ 2000 (Invitrogen). siRNA transfections were performed with Lipofectamine RNAiMAX (Invitrogen).

293FT or HepG2 cells expressing IR-GFP WT, or mutants were generated as described previously^28^. Briefly, cDNAs encoding IR mutants were cloned into the pBabe-GFP-puro vector. The vectors were co-transfected with viral packaging vectors into 293FT cells, and the viral supernatants were collected at 2 days and 3 days after transfection. The concentrated viruses were added to 293FT and HepG2 cells with 4 µg/ml of polybrene. Cells were selected with puromycin (1 µg/ml for 293FT and 2 µg/ml for HepG2) at 3 days after infection and sorted by flow cytometry to collect cells expressing similar levels of IR-GFP.

For the expression of IRS1 WT or mutants in HepG2 cells, cDNA encoding IRS1 WT or mutants were cloned into the pBabe-mCherry-puro vector. The vectors were co-transfected with viral packaging vectors into 293FT cells, and the viral supernatants were collected. The concentrated viruses were added to HepG2 cells stably expressing IR-GFP WT. The cells were transfected with the indicated siRNA at 1 day after viral infection. Analysis was performed at 4 days after infection.

Primary hepatocytes were isolated from 2-month-old male mice with a standard two-step collagenase perfusion procedure. Cells were plated on collagen-coated dishes and incubated in attachment medium [Williams’ E Medium supplemented with 5% (v/v) FBS, 10 nM insulin, 10 nM dexamethasone, and 1% (v/v) penicillin/streptomycin]. After 2-4 h, the medium was changed to low-glucose DMEM supplemented with 5% (v/v) FBS, 10 nM dexamethasone, 10 nM insulin, 100 nM triiodothyronine, and 1% (v/v) penicillin/streptomycin. After 1 day, the cells were serum starved for 14 h and treated with dimethyl sulfoxide (DMSO) or SHP099 for 2 h.

Adeno-associated viruses (AAV) encoding SHP2 shRNA (AAV8-GFP-U6-mPTPN11-shRNA) or control shRNA (AAV-GFP-U6-scrmb-shRNA) were generated at Vector Biolabs. We injected 10-11-week-old male mice fed with HFD for 5 weeks with the viruses at 1 × 10^12^ genomic copies per mouse. Experiments were performed at 2 weeks after virus injection.

### Tissue histology and immunohistochemistry

The fixation, histological analysis, and immunohistochemistry of mouse tissues were performed as described previously^28^. For human patient sample analysis, the deparaffinized sections were subjected to antigen retrieval with 10 mM sodium citrate (pH 6.0), incubated with 0.3% H_2_O_2_, blocked with 0.3% BSA, and then incubated first with anti-IR (CT3, Millipore, 1:100) and anti-ZO1 (Thermo Scientific, 1:200) antibodies and then with secondary antibodies (Alexa Fluor 568 goat anti-mouse and Alex Fluor 488 goat anti-rabbit; Molecular Probes). The slides were counterstained with DAPI. Five to nine images (depends on the percentage of normal hepatocytes) were randomly taken under 40x magnification. The total cell numbers and numbers of IR PM-positive cells were counted at least twice for individual images. Over 100 cells were analyzed for each patient samples. All immunohistochemistry and scoring were performed blinded to the diabetes status. Human tissue collection and analysis were performed under the supervision of the Institutional Review Board (IRB; STU 062013-043).

### Metabolic profiling

Blood samples were collected from overnight fasted mice using submandibular bleeding methods. For serum preparation, blood was allowed to form clots at room temperature for 30 min, centrifuged at 3000 g for 15 min at 4°C, and stored at −80°C. Serum insulin and C-peptide levels were determined with the STELLUX Chemic Rodent Insulin ELISA kit (Alpco) and Mouse C-peptide ELISA kit (Alpco), respectively. ELISA analysis for insulin and C-peptide was performed by Metabolic Phenotyping Core at UT Southwestern Medical Center.

### Immunoprecipitation (IP) and quantitative Western blots

Cells were incubated with the cell lysis buffer [50 mM HEPES (pH 7.4), 150 mM NaCl, 10% (v/v) Glycerol, 1% (v/v) Triton X-100, 1 mM EDTA, 100 mM sodium fluoride, 2 mM sodium orthovanadate, 20 mM sodium pyrophosphate, 0.5 mM dithiothreitol (DTT), 2 mM phenylmethylsulfonyl fluoride (PMSF)] supplemented with protease inhibitors (Roche) and PhosSTOP (Roche) on ice for 1 h. The cell lysates were cleared by centrifugation and incubated with antibody-conjugated beads. The beads were washed, and the bound proteins were eluted with the SDS sample buffer and analyzed by SDS-PAGE and Western blotting. For quantitative Western blots, anti-rabbit immunoglobulin G (IgG) (H+L) (Dylight 800 or 680 conjugates) and anti-mouse IgG (H+L) (Dylight 800 or 680 conjugates) (Cell Signaling) were used as secondary antibodies. The membranes were scanned with the Odyssey Infrared Imaging System (LI-COR, Lincoln, NE).

### Immunofluorescence

Indirect immunofluorescence microscopy was performed on cells grown on coverslips and fixed in cold methanol at −20°C for 10 min. The fixed cells were incubated with PBS for 30 min and 3% BSA in 0.1% PBST for 1 h, and then treated with diluted antibodies in 0.3% BSA in 0.1% PBST at 4°C overnight. After being washed, cells were incubated with fluorescent secondary antibodies and mounted on microscope slides in ProLong Gold Antifade reagent with DAPI (Invitrogen). Images of fixed cells were acquired as a series of 0.4 µm stacks with a DeltaVision system (Applied Precision, Issaquah, WA). Raw images were deconvolved using the iterative algorithm implemented in the softWoRx software (Applied precision, Issaquash, WA). The central section of a 0.4 µm z-stack containing 3 contiguous focal planes was used for quantification. The cell edges were defined with Image J. The whole cell signal intensity (WC) and intracellular signal intensity (IC) were measured. The plasma membrane signal intensity (PM) was calculated by subtracting IC from WC. Identical exposure times and magnifications were used for all comparative analyses.

### Protein purification

The full-length human AP2M1 was cloned into a pGEX 6P-1, and the plasmid was transformed into *Escherichia coli* strain BL21 (DE3). Protein expression was induced by 0.2 mM isopropyl β-D-1-thiogalactopyranoside (IPTG) at 25°C overnight. The harvested pellets were lysed in the lysis buffer I [20 mM Tris-HCl, pH 8.0, 150 mM NaCl, 1% (v/v) TritonX-100, 5% (v/v) Glycerol, 1 mM DTT, 1 mM PMSF]. After sonication, lysates were cleared by centrifugation at 4°C. The supernatants were filtered by 0.45 µm filter and incubated with pre-equilibrated Glutathione Sepharose 4B beads (GE Healthcare). The resulting protein-bound beads were washed extensively with lysis buffer I.

The AP2M1 fragment (residues 160-435) was cloned into a modified pET28a that introduced an N-terminal His_6_-tag followed by a thrombin cleavage site. The plasmid was transformed into BL21 (DE3). Protein expression was induced by 0.2 mM IPTG at 20°C overnight. The harvested pellets were lysed in the lysis buffer II [20 mM Tris-HCl, pH 7.5, 500 mM NaCl, 20 mM Imidazole, 1 mM PMSF]. After sonication, lysates were cleared by centrifugation at 4°C. The supernatants were filtered by 0.45 µm filter and incubated with pre-equilibrated Ni2^+^-NTA beads (Qiagen). Protein-bound beads were washed with 150 ml of wash buffer I [20 mM Tris-HCl, pH 7.5, 1M NaCl, 20 mM Imidazole] and with 50 ml of wash buffer II [20 mM Tris-HCl, pH 7.5, 100 mM NaCl, 20 mM Imidazole]. The proteins were then eluted with the elution buffer [20 mM Tris-HCl, pH 7.5, 100 mM NaCl, 150 mM Imidazole] and incubated with thrombin (Sigma) at 4°C overnight. The protein was further purified with a Superdex 200 size exclusion column (GE Healthcare). The relevant protein fractions were pooled, aliquoted, and snap-frozen for future experiments.

### Crystallization of the AP2M1-pIRS1 complex

The purified AP2M1 (residues 160-435) was mixed with the pS-IRS1 peptide (CHTDDGYMPMpSPGVA, residues 607-620) at a molar ratio of 1:5 and then crystalized with the hanging-drop vapor diffusion method. The crystals of the AP2M1-pS-IRS1 complex grew within few days after the protein solution was mixed with the reservoir solution [1.0 M sodium malonate, pH 5.0, 0.1 M sodium acetate tri-hydrate, pH 4.5, 2% (w/v) PEG 20k]. All crystals were cryoprotected with the reservoir solution including 15% (w/v) glycerol for data collection.

### Data collection and structure determination

X-ray diffraction datasets were collected at the Advanced Photon Source (APS) beamline Sector 19-ID at a wavelength of 0.97914 Å and at 100K. HKL3000 was used to process the datasets^72^. The crystal of the AP2M1-pS-IRS1 complex diffracted to a minimum Bragg spacing of 3.2 Å and exhibited the symmetry of space group P6_4_ with cell dimensions of a = b = 125.33 Å, c = 74.82 Å. There are two molecules in the asymmetric unit, with a 53.4% solvent content.

The structure was determined by molecular replacement with PHASER-MR ^73^, using the structure of the AP2M-IGN38 complex (PDB ID: 1BXX) as the search model. Structure refinement was performed with COOT and PHENIX^74-76^. The final R_work_ and R_free_ were 20.3% and 23.6%, respectively. Data collection and refinement statistics are provided in Supplementary Table 1. The model quality was validated with Molprobity^77^. All structural figures were generated with the program PyMOL (http://www.pymol.org/) with the same color and labeling schemes.

### Protein-binding assays

For GST pull-down assays of in vitro translated (IVT) IRS1 or IRS2 proteins, beads bound to GST-AP2M1 or GST were incubated with IVT products diluted in the cell lysis buffer at 4°C for 2 h. After incubation and washing, proteins bound to beads were eluted with the SDS loading buffer, resolved with SDS-PAGE, and detected with Coomassie staining or immunoblotting with the appropriate antibodies. Peptide pull-down assays were performed as described previously^28^. The isothermal titration calorimetry (ITC) assays were performed with a MicroCal Omega ITC200 titration calorimeter (GE Life Sciences) at 20°C with minor modifications^78^. Briefly, the recombinant AP2M1 protein (residues 160-435) and peptides were dialyzed into the HEPES buffer [25 mM HEPES, pH 7.5, 50 mM NaCl]. For each titration, 300 µl of AP2M1 (50 µM) were added to the calorimeter cell. IRS1 peptides (YMPMS, 528.9 µM or YMPMpS, 507.4µM) were injected with an injection syringe in nineteen 2.0-µl portions. Raw data were processed and fitted with the NITPIC software package ^79^.

### Phosphatase assays

Active SHP2 (2.9 µM, SignalChem) diluted in the phosphatase dilution buffer [50 mM imidazole, pH 7.2, 0.2% 2-mercaptoethanol, 65 ng/µl BSA] was incubated with IRS1 peptides (2.6 mM) at 37°C for the indicated time points. Two microliters of reaction products were spotted onto 0.45 µm nitrocellulose membrane (BioRad) and dried completely. The membrane was blocked with 5% nonfat milk in TBS for 1h, and washed once with TBS-T (0.02% Tween 20). The membrane was incubated with anti-IRS1-pY612 antibodies diluted in TBS-T at 4°C overnight. After washing with TBS-T, the anti-rabbit immunoglobulin G (IgG) (H+L) Dylight 800 conjugates (Cell Signaling) were applied as secondary antibodies. The membranes were scanned with the Odyssey Infrared Imaging System (LI-COR, Lincoln, NE) for quantification.

### In vivo insulin signaling assays

HFD-fed mice were administered vehicle or SHP099 for 5 days. The mice were fasted overnight and administered vehicle or SHP099 once more. At 2 h after the last administration, the mice were injected with or without 1 U insulin (Eli Lilly) via inferior vena cava. For hepatic knockdown of SHP2, mice fed HFD were injected with AAV-GFP or SHP shRNA. At 17 days after injection, the mice were fasted overnight and injected with or without 1 U insulin (Eli Lilly) via inferior vena cava. The livers were collected at the indicated time points. White adipose tissue (WAT) and skeletal muscles were collected at 2 min and 3 min after the indicated time points, respectively. 30 mg of tissue was mixed with the cell lysis buffer [50 mM HEPES (pH 7.4), 150 mM NaCl, 10% (v/v) glycerol, 1% (v/v) Triton X-100, 1 mM EDTA, 100 mM sodium fluoride, 2 mM sodium orthovanadate, 20 mM sodium pyrophosphate, 0.5 mM dithiothreitol (DTT), 2 mM phenylmethylsulfonyl fluoride (PMSF)] supplemented with protease inhibitors (Roche), PhosSTOP (Roche), 25 U/ml turbo nuclease (Accelagen) and 100 µM cytochalasin B (Sigma), homogenized with Minilys (Bertin Technologies), and then incubated on ice for 1 h. After centrifugation at 20,817 g at 4°C for 1 h, the lysates were analyzed by quantitative Western blotting.

### Statistical analyses

Prism was used for the generation of all curves and graphs and for statistical analyses. Results are presented as mean ± SEM or mean ± SD. Two-tailed unpaired t tests were used for pairwise significance analysis. Sample sizes were determined on the basis of the maximum number of mice that could be bred in similar ages to maintain well-matched controls. Power calculations for sample sizes were not performed. We monitored weight and health conditions of mice, and excluded mice from experiments if the animal was unhealthy and the body weight was more than two standard deviations from the mean. Randomization and blinding methods were not used, and data were analyzed after the completion of all data collection in each experiment.

## Data availability

The coordinates of the AP2M1-pIRS1 structure have been deposited to the Protein Data Bank (PDB ID: 6BNT). All other data are available from the authors upon request.

## Acknowledgements

We thank Drs. Joseph Goldstein, Michael Brown, David Mangelsdorf, and Philipp Scherer for advice on animal experiments and helpful discussions, Dr. Melanie Cobb for advice on MEK inhibitors and helpful discussions, Dr. Sandra Schmid for advice on IR endocytosis and key reagents, Dr. Zhonghui Lin for advice on structural analysis, Drs. James A. Richardson and John Shelton for advice on histopathology, and Dr. Xuelian Luo for advice on protein purification and helpful discussions. We are grateful to the Animal Resource Center at University of Texas Southwestern Medical Center for assistance with mouse maintenance, Metabolic Phenotyping Core for assistance with analysis of insulin and C-peptide, and the Histopathology Core for assistance with tissue processing and sectioning. This work is supported by the Clayton Foundation and the National Institutes of Health (1R01GM124096). H.Y. is an Investigator with the Howard Hughes Medical Institute.

## Author contributions

E.C. and H.Y. designed the study and wrote the paper. E.C. performed all animal experiments, cellular experiments, human patient sample analysis and most biochemical experiments. S.K. determined and analyzed the structure of AP2M1 bound to the phospho-IRS1 peptide. H.G. conducted the isothermal titration calorimetry (ITC) assay. K.B. generated HepG2 cell lines expressing IR mutants and performed cellular experiments. I.N., A.Y., A.S., and H.Z. provided human liver samples and expertise on analysis.

## Supplemental Figures

**Supplementary Fig. 1.**
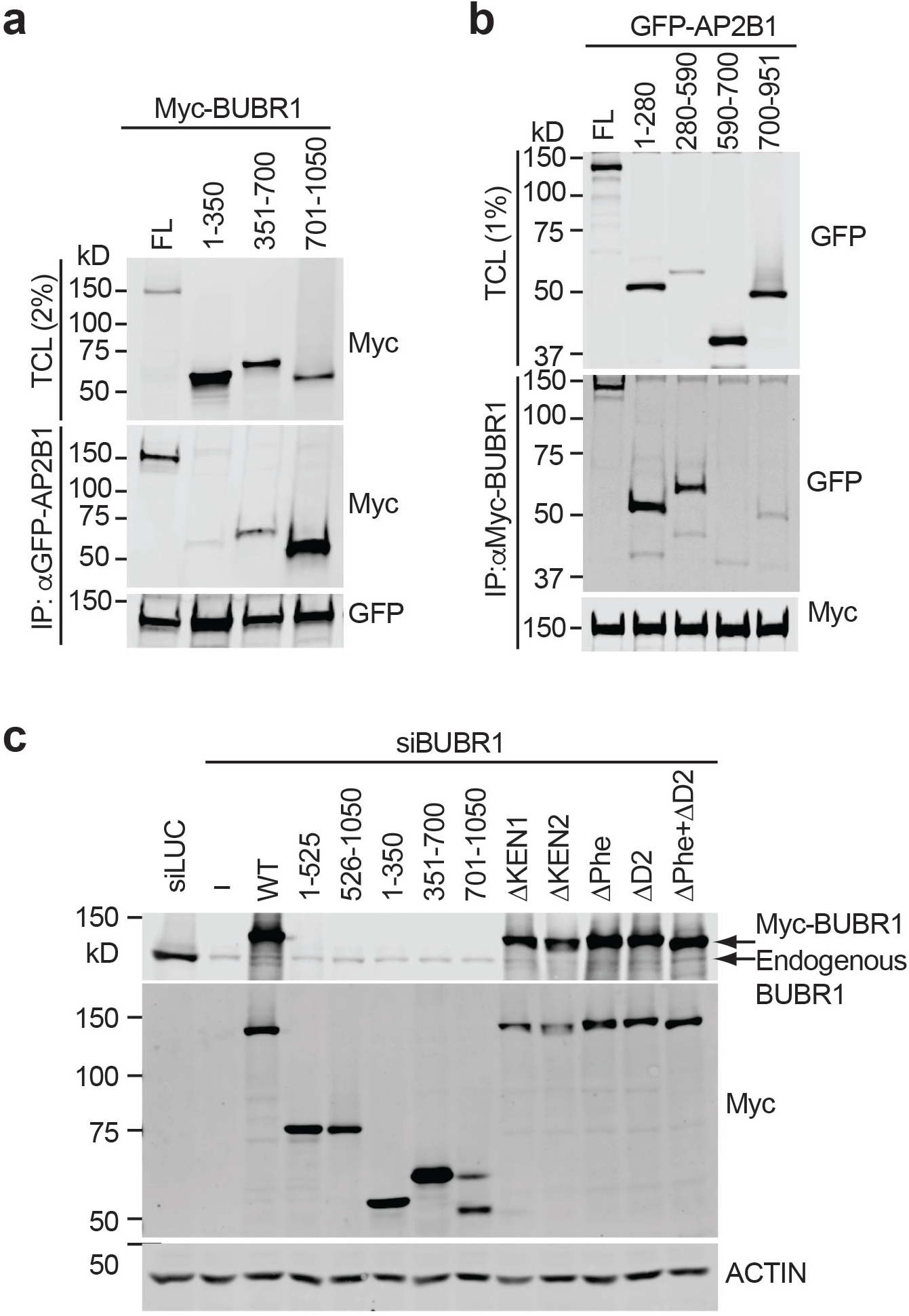
Mapping the binding domains between BUBR1 and AP2B1. **a** Binding of BUBR1 full length (FL) and truncation mutants to AP2B1. 293FT cells were transfected with plasmids encoding GFP-AP2B1 and Myc-BUBR1 proteins. Total cell lysate (TCL) and anti-GFP-AP2B1 IP were blotted with anti-Myc and anti-GFP antibodies. **b** Binding of AP2B1 full length (FL) and truncation mutants to BUBR1. 293FT cells were transfected with plasmids encoding GFP-AP2B1 and Myc-BUBR1 proteins. Total cell lysate (TCL) and anti-Myc-BUBR1 IP were blotted with anti-GFP and anti-Myc antibodies. **c** Western blot analysis of lysates of cells in Fig. 1h.

**Supplementary Fig. 2.**
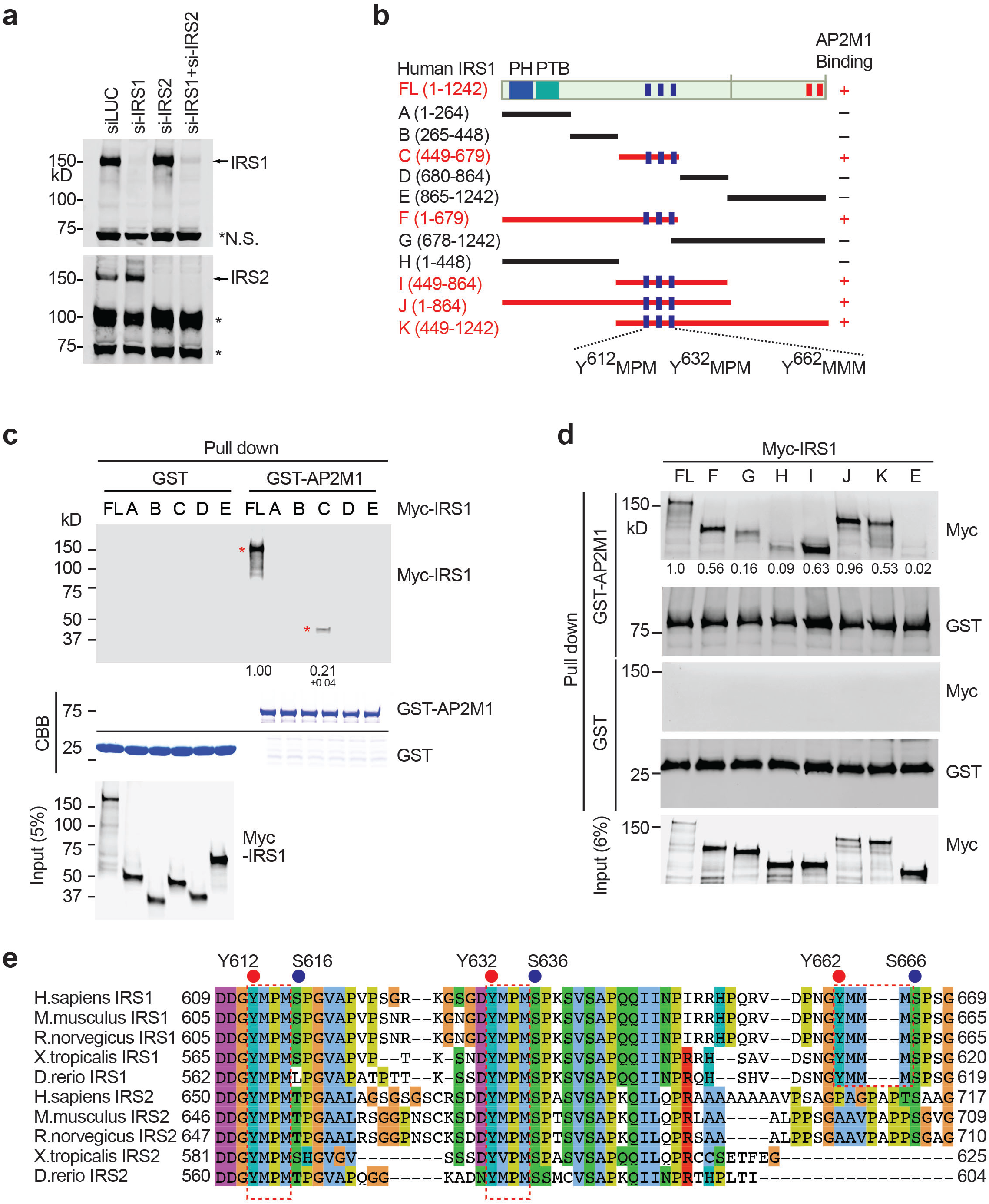
IRS1 promotes IR endocytosis and interacts with AP2. **a** Western blot analysis of cell lysates in Fig. 3a. Asterisks indicate non-specific bands. **b** Domains and YXXΦ motifs of human IRS1. PH, pleckstrin homology domain; PTB, phosphotyrosine-binding domain. IRS1 fragments that can or cannot bind to AP2M1 are presented as red or black lines, respectively. YXXΦ motifs and phosphotyrosine sites for SHP2 binding are presented as blue and red bars, respectively. **c** Binding of IRS1 WT and mutants to GST or GST-AP2M1. Input and protein bound to beads were blotted with anti-Myc (IRS1) antibodies and stained with Coomassie (CBB). The relative band intensities are shown below (mean ± SD; n=3 independent experiments). **d** Binding of IRS1 WT and truncation mutants to GST or GST-AP2M1. Input and protein bound to beads were blotted with the indicated antibodies. The relative band intensities are shown below (n=2 independent experiments). **e** Sequence alignment of a conserved region in IRS1/2. Three YXXΦ motifs are boxed with red dashed lines. The phosphorylation sites of IR and MAPK are indicated as red and blue dots, respectively.

**Supplementary Fig. 3.**
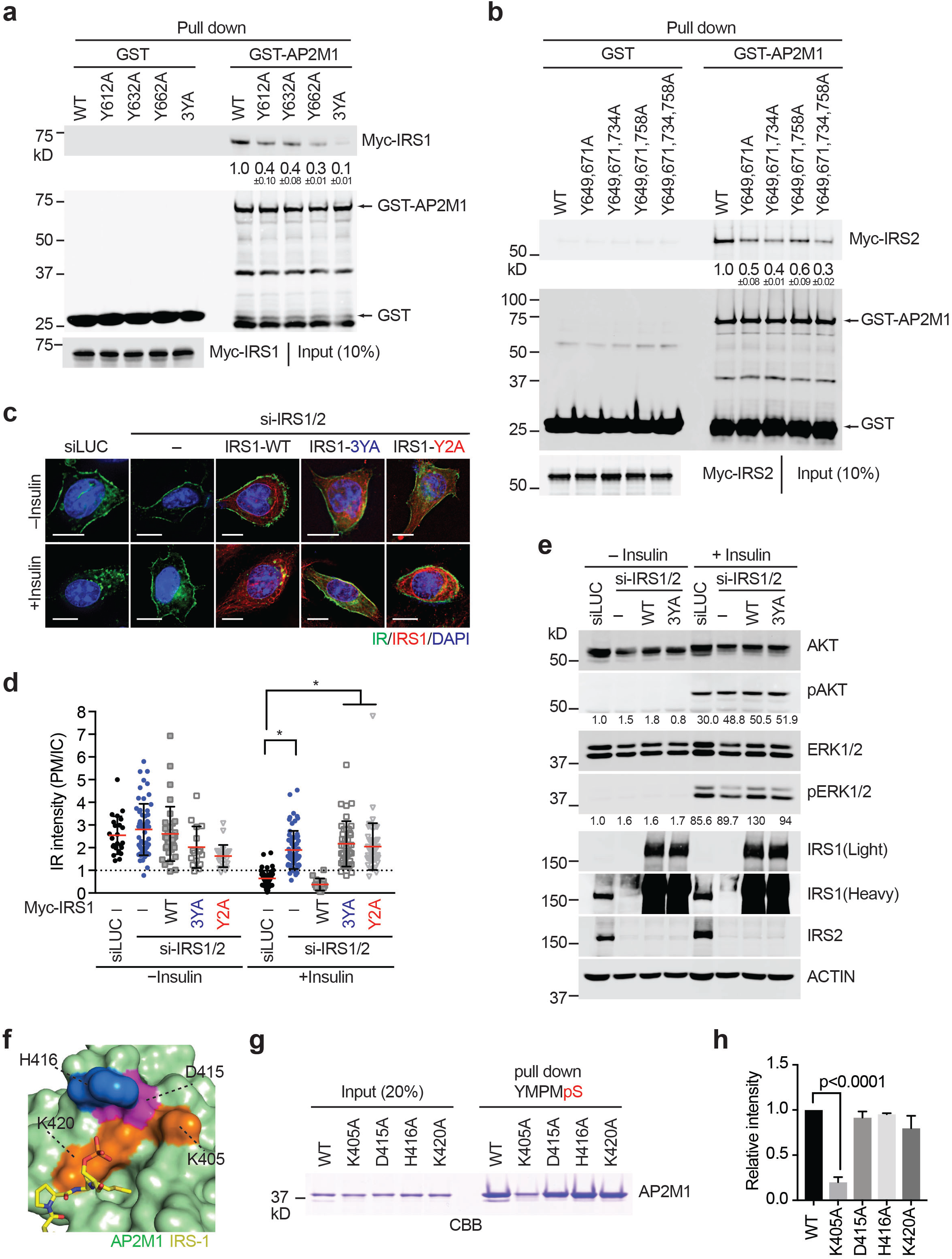
The YXXΦ motifs of IRS1/2 bind to AP2M1 and are required for insulin-activated IR endocytosis. **a** Binding of IRS1 WT and mutants to GST or GST-AP2M1. The relative band intensities are shown below (3YA, Y612A/Y632A/Y662A; mean ± SD; n=3 independent experiments). **b** Binding of IRS2 WT and mutants to GST or GST-AP2M1. The relative band intensities are shown below (Mean ± SD; n=2 independent experiments). **c** HepG2 cells stably expressing IR-GFP WT were transfected with the indicated siRNA or siRNA-resistant mCherry-IRS1, serum starved, treated without or with 100 nM insulin for 5 min, and stained with anti-GFP (IR; green), anti-mCherry (IRS1; red), and DAPI (blue). (3YA, Y612A/Y632A/Y662A; Y2A, Y1179A/Y1229A). **d** Quantification of the ratios of PM and IC IR-GFP signals of cells in (**c**) (mean ± SD; *p<0.0001). **e** 293FT cells stably expressing IR-GFP WT were transfected with the indicated siRNA or siRNA-resistant Myc-IRS1, serum starved, treated without or with 100 nM insulin form 5min. The cell lysates were subjected to quantitative immunoblotting with the indicated antibodies. The relative band intensities are shown below. Representative results from 3 independent experiments were presented. **f** Surface drawing of AP2M1 with the bound pS-IRS1 shown as sticks. The potential acceptor residues for IRS1 pS616 are labeled. **g** Binding of the pS-IRS1 peptide to WT and mutants of AP2M1 (residues 160-435). Input and proteins bound to pS-IRS1 peptides were analyzed by SDS-PAGE and stained with Coomassie (CBB). **h** Quantification of the relative band intensities in (**g**). Mean ± SD; n=3 independent experiments.

**Supplementary Fig. 4.**
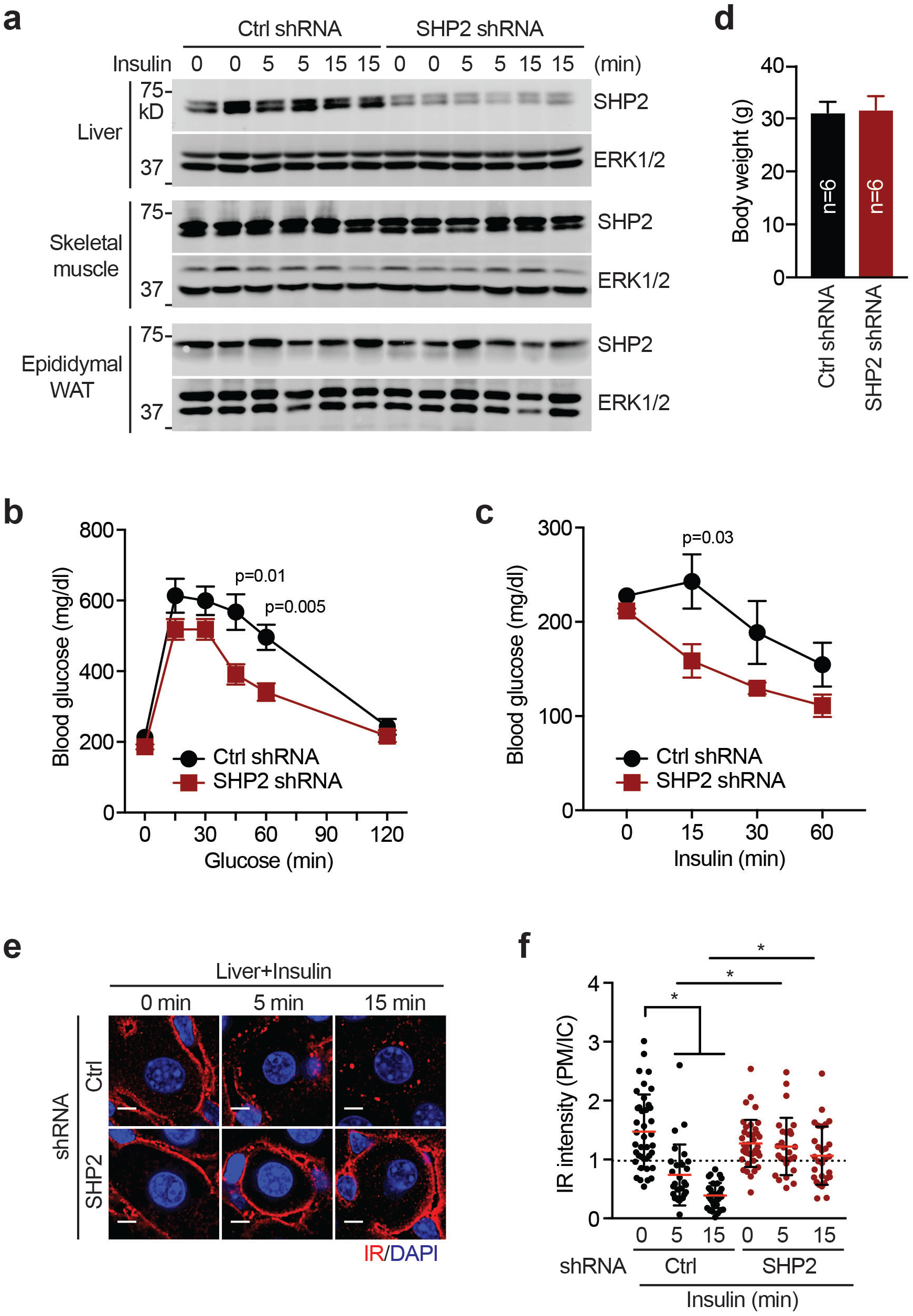
Depletion of SHP2 by shRNA delays IR endocytosis and improves insulin sensitivity in mice. **a** The level of SHP2 in liver, skeletal muscle and epididymal WAT from mice fed HFD for 5 weeks. The mice were injected with AAV-control (Ctrl) or SHP2 shRNA. At 17 days after injection, the mice were fasted overnight and injected with or without 1 U insulin via inferior vena cava. The livers were collected at the indicated time points. WAT and skeletal muscle were collected at 2 min and 3 min after the indicated time points, respectively. Lysates were prepared from these tissues and subjected to quantitative immunoblotting with the indicated antibodies. **b**,**c** Glucose tolerance test (**b**) and insulin tolerance test (**c**) in mice injected with AAV-Ctrl or AAV-SHP2 shRNA and fed HFD. Experiments were performed at 2 weeks after injection. n=6; mean ± SEM. **d** Body weight in HFD-fed mice injected with AAV-Ctrl or AAV-SHP2 shRNA. Mean ± SD. **e** HFD-fed mice were injected with AAV-Ctrl or AAV-SHP2. At 17 days after injection, the mice were fasted overnight and injected with or without 1U insulin via inferior vena cava. The livers were collected at the indicated time points and the sections were stained with anti-IR (red) and DAPI (blue). Scale bars, 5 µm. **f** Quantification of the ratios of PM and IC IR signals of the livers in (**e**) (mean ± SD; *p<0.0001).

**Supplementary Fig. 5.**
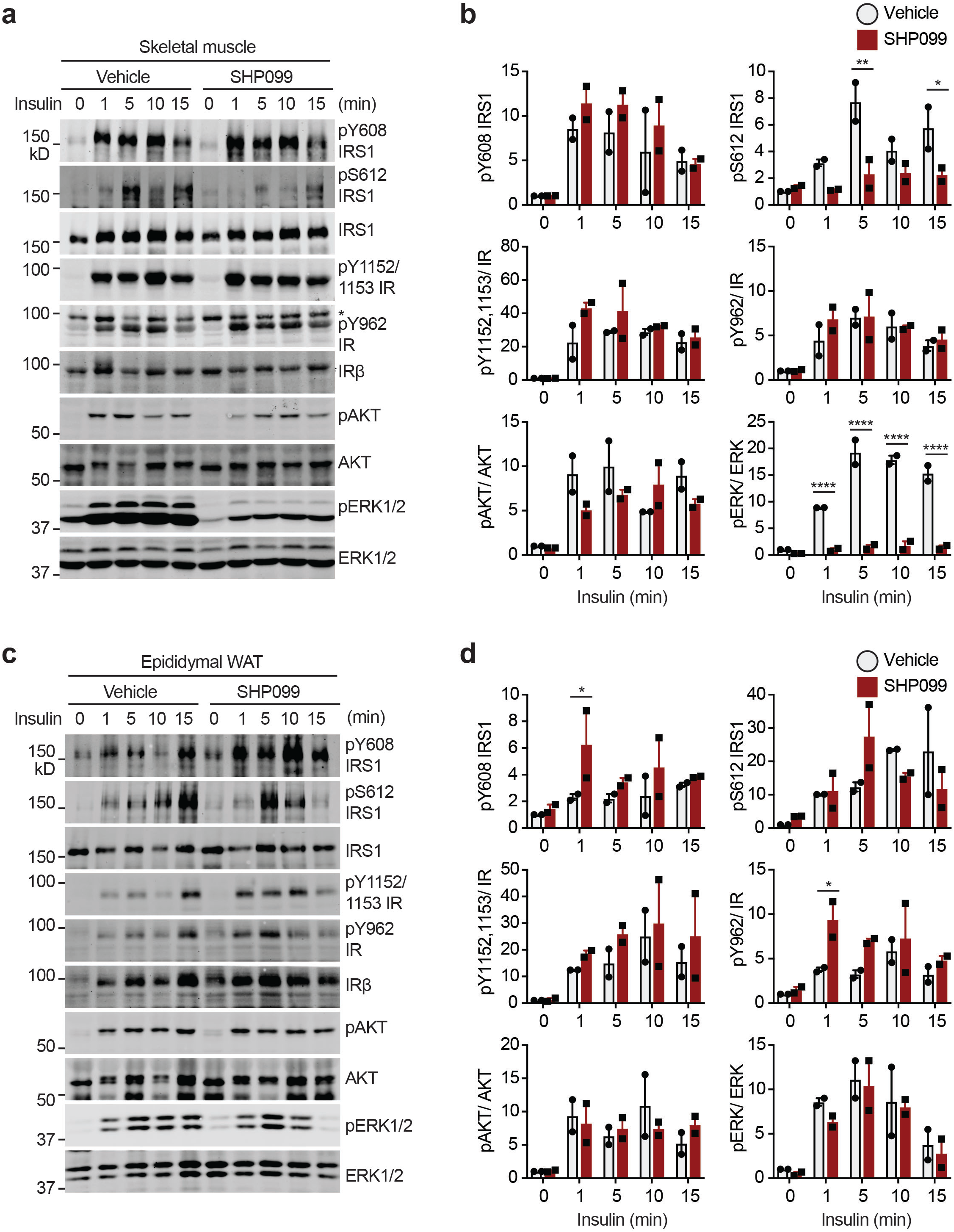
Effect of SHP099 on insulin signaling in skeletal muscle and white adipose tissue (WAT) from HFD-fed mice. **a** Insulin signaling in the skeletal muscle from mice fed HFD for 5 weeks. The mice were administered vehicle or SHP099 for 5 days, fasted overnight and administered vehicle or SHP099 once more. At 2 h after the last administration, the mice were injected with or without 1 U insulin via inferior vena cava. The livers were collected at the indicated time points (See Fig. 6a,b). WAT and skeletal muscle were collected at 2 min and 3 min after the indicated time points, respectively. Lysates were prepared from these tissues and subjected to quantitative immunoblotting with the indicated antibodies. Asterisk indicates non-specific bands. **b** Quantification of the blots in (**a**). Mean ± SD; *p<0.05, **p<0.01, ***p<0.001, and ****p<0.0001. **c** Insulin signaling in the WAT from mice in (**a**). **d** Quantification of the blots in (**c**). Mean ± SD; *p<0.05.

**Supplementary Table 1.** Data processing and refinement statistics.

|  | AP2M1-pIRS1 |
| --- | --- |
| **Data collection** |  |
| Space group | P6<sub>4</sub> |
| Cell dimensions |  |
| *a*, *b*, *c* (Å) | 125.33, 125.33, 74.82 |
| $\alpha$, $\beta$, $\gamma$ (°) | 90.00, 90.00, 120.00 |
| Resolution (Å) | 32.6–3.20 (3.26–3.20) |
| *R*<sub>merge</sub> (%) | 16.1 (29.2) |
| $\langle I \rangle / \langle \sigma_I \rangle$ | 13.2 (1.4) |
| Completeness (%) | 100 (100) |
| Number of total reflections | 152046 |
| Number of unique reflections | 11485 |
| Redundancy | 12.4 (13.3) |
| **Refinement** |  |
| Resolution (Å) | 27.14–3.20 (3.31–3.20) |
| No. reflections (work/free) | 11157 (1113) / 1113 (113) |
| *R*<sub>work</sub> / *R*<sub>free</sub> | 20.3 (29.1) / 23.6 (35.9) |
| R.m.s deviations |  |
| Bond lengths (Å) | 0.01 |
| Bond angles (°) | 1.30 |
| Completeness (%) | 100 |
| Ramachandran plot |  |
| Favored (%) | 91 |
| Allowed (%) | 8.4 |
| Outliers (%) | 0.4 |

\*Highest-resolution shell is shown in parenthesis.

